# Multiple plastid losses within photosynthetic stramenopiles revealed by comprehensive phylogenomics

**DOI:** 10.1101/2024.02.03.578753

**Authors:** Kristina X. Terpis, Eric D. Salomaki, Dovilė Barcytė, Tomáš Pánek, Heroen Verbruggen, Martin Kolisko, J. Craig Bailey, Marek Eliáš, Christopher E Lane

**Affiliations:** Department of Biological Sciences, University of Rhode Island, Kingston RI, USA; Biology Centre, Czech Academy of Sciences, České Budějovice, Czech Republic; Center for Computational Biology of Human Disease and Center for Computation and Visualization, Brown University, Providence, RI, USA; Department of Biology and Ecology, Faculty of Science, University of Ostrava, Czech Republic; Department of Zoology, Faculty of Science, Charles University, Prague, Czech Republic; School of BioSciences, University of Melbourne, Victoria 3010, Australia; Department of Biology and Marine Biology, University of North Carolina, Wilmington, NC, USA

**Keywords:** Ochrophyta, phylogenomics, *Picophagus flagellatus*, plastid loss, Stramenopiles, Synchromophyceae

## Abstract

Ochrophyta is a vast and morphologically diverse group of algae with complex plastids, including familiar taxa with fundamental ecological importance (diatoms or kelp), and a wealth of lesser-known and obscure organisms. The sheer diversity of ochrophytes poses a challenge for reconstructing their phylogeny, with major gaps in sampling and an unsettled placement of particular taxa yet to be tackled. We sequenced transcriptomes from 25 strategically selected representatives and used these data to build the most taxonomically comprehensive ochrophyte-centered phylogenomic supermatrix to date. We employed a combination of approaches to reconstruct and critically evaluate the relationships among ochrophytes. While generally congruent with previous analyses, the updated ochrophyte phylogenomic tree resolved the position of several taxa with previously uncertain placement, and supported a redefinition of the class Synchromophyceae. Our results indicated that the heterotrophic plastid-lacking heliozoan *Actinophrys sol* is not a sister lineage of ochrophytes, as proposed recently, but rather phylogenetically nested among them. In addition, we found *Picophagus flagellatus* to be a secondarily heterotrophic ochrophyte lacking all hallmark plastid genes, yet exhibiting mitochondrial proteins that seem to be genetic footprints of lost plastid organelle. We thus document, for the first time, plastid loss in two separate ochrophyte lineages. Altogether, our study provides a new framework for reconstructing trait evolution in ochrophytes and demonstrates that plastid loss is more common than previously thought.

**Issue Section:** Discoveries

## Introduction

The stramenopiles, alveolates, and rhizarians form one of the most diverse major eukaryotic supergroups, commonly referred to as SAR (Burki et al. 2007; Grattepanche et al. 2018). Some members of SAR have been intensely studied, particularly parasites including oomycetes (stramenopiles), which cause millions of dollars in damages to crops annually, or apicomplexans, like *Plasmodium* (alveolates), which is the causative agent of malaria. Photosynthetic lineages, such as diatoms and kelp (stramenopiles), and dinoflagellates (alveolates) have also been extensively studied due to their importance as primary producers in marine ecosystems. However, many members of this supergroup have received far less attention (Massana et al. 2004; Lin et al. 2012; Grattepanche et al. 2018). Stramenopiles, also called heterokonts, stem from a biflagellated ancestor with the anterior flagellum covered by characteristic tripartite mastigonemes, although this stramenopile synapomorphy has been variously modified in many extant lineages (Thakur et al. 2019). Surveys of marine environments have revealed that stramenopiles make up a large fraction of the unstudied microscopic marine eukaryotes (Massana et al. 2004; Logares et al. 2014; Pernice et al. 2016; Obiol et al. 2023).

Photosynthesis has a complex evolutionary history across SAR, exemplified by multiple independent plastid acquisitions across the supergroup (Delwiche 1999; Keeling 2013; Derelle et al. 2016; Ševčíková et al. 2016; Dorrell et al. 2017; Grattepanche et al. 2018). Stramenopiles themselves include a single photosynthetic clade, termed Ochrophyta (Cavalier-Smith et al. 1995). The ochrophytes include members ranging in complexity from unicellular diatoms that are critical for primary production in the ocean, to 30m multicellular kelp, which are important ecosystem builders. They are found in marine, freshwater, and terrestrial habitats and exhibit a wide range of morphologies, life histories, and nutritional strategies. Ochrophytes have a plastid of red algal origin, acquired via an endosymbiotic event with the common ancestor of ochrophytes (Stiller et al. 2014; Ševčíková et al. 2016; Dorrell et al. 2017). Most ochrophyte plastids have chlorophylls *a* and *c*, with fucoxanthin as the predominant xanthophyll (Andersen 2004). However, photosynthetic pigment profiles vary considerably, and while generally considered diagnostic for different ochrophyte subgroups, parallel losses of accessory pigments have led to the incorrect assignment of some species among ochrophyte classes (Ott et al. 2015).

Even though photosynthesis is the dominant source of energy among ochrophytes, many are mixotrophs, and several independent lineages have lost all photosynthetic pigments and became secondarily heterotrophic (Grant et al. 2009; Kamikawa et al. 2017; Dorrell et al. 2019; Onyshchenko et al. 2019). All non-photosynthetic ochrophytes investigated in sufficient detail proved to have retained a residual plastid (Barcytė et al. 2024), but some of them, such as the tiny marine colorless flagellate *Picophagus flagellatus* (Guillou et al. 1999), remain unstudied in this regard. An interesting case is the heterotrophic actinophryids – a lineage originally placed within Heliozoa, but determined to have ultrastructural affinities for stramenopiles (Patterson 1979; Patterson 1986). Molecular evidence later confirmed this assessment as a stramenopile subgroup, with phylogenetic affinities to ochrophytes (Nikolaev et al. 2004; Cavalier-Smith and Scoble 2013). The first phylogenomic study of *Actinophrys sol* inferred it as a sister lineage to all ochrophytes and revealed that it lacks a plastid, yet contains potential genomic footprints of a past photosynthetic endosymbiont (Azuma et al. 2022).

The evolutionary relationships among ochrophytes were first hypothesized based upon morphological traits, however, the use of molecular markers for phylogenetics (particularly the nuclear 18S rRNA and the plastidial *rbcL* genes) identified species that were ultimately described as new classes, which could not be distinguished by morphology alone (e.g., Bailey et al. 1998; Kawachi et al. 2002). The first multi-gene data used to resolve the evolutionary relationships among ochrophytes employed a five-gene alignment, recovering three main clades termed SI, SII, and SIII (Yang et al. 2012). Shortly after, the Marine Microbial Eukaryotic Sequencing Project (MMETSP) represented a fundamental turning point in data availability for stramenopiles. Over 650 novel transcriptomes were sequenced from across the tree of life (Keeling et al. 2014) and nearly half of the data generated (269 transcriptomes) were from stramenopiles. This effort was intentional, due to the low amount of stramenopile data relative to the diversity of the group. However, even the MMETSP was heavily biased towards diatoms (Bacillariophyceae *sensu lato*). Despite taxonomic biases, these data were combined with additional new transcriptomes to produce the first phylogenomic analysis of ochrophytes (Derelle et al. 2016). While there was robust support for the monophyly and relationships among the classes represented, only data from nine out of 18 classes recognized today were included. Subsequently, the expanded application of high-throughput sequencing enabled phylogenomic studies to significantly improve the resolution of relationships within and among ochrophyte classes and stramenopile lineages overall (Noguchi et al. 2016; Jackson et al. 2017; Leonard et al. 2018; Dorrell et al. 2019; Dorrell et al. 2021; Cho et al. 2022).

The range of morphology, life history, ecology, and overall diversity of ochrophytes provides enormous scope for studying the evolution of these organismal traits. However, the lack of a resolved ochrophyte phylogeny, representing the diversity of the group, significantly hinders the ability to make meaningful inferences about their evolution. To improve the resolution of the ochrophyte phylogeny, we inferred a densely sampled ochrophyte phylogeny from nuclear-encoded genes, with our sampling design aimed at improving class-level representation, placing historically difficult to resolve taxa and identifying cases of plastid loss. Furthermore, by reevaluating the phylogenetic position of the actinophryid *A. sol* and examining the genome of *Picophagus flagellatus* for the genetic building blocks of a plastid, we identified two distinct extant phagotrophic free-living lineages within Ochrophyta that have completely lost their plastids.

## Results and Discussion

### A new highly resolved phylogeny of ochrophytes

In an effort to clarify the evolutionary relationships among ochrophytes, we generated transcriptomes from 25 ochrophytes. These taxa were chosen from lineages with little to no publicly available high-throughput sequencing data, including those considered *incertae sedis* within Ochrophyta. The taxonomic identity of the targeted organisms was checked and updated according to the most recent taxonomic revisions, so the names we use may differ from those in the culture collections (table S1). We combined our data with other publicly available genomic and transcriptomic data, including those not previously utilized for phylogenomics (Derelle et al. 2016; Azuma et al. 2022; Di Franco et al. 2022; Barcytė et al. 2024; Agostini et al. in prep.), to build the most taxonomically comprehensive phylogenomic dataset for stramenopiles to date, including nearly all major ochrophyte lineages defined by an 18S rRNA gene phylogeny that have representatives available in culture (fig. S1).

Using the recently developed software package PhyloFisher (Tice et al. 2021; Jones et al. 2024), we created two datasets to explore evolutionary relationships 1) among stramenopiles, and 2) specifically among ochrophytes. The more comprehensive of these datasets (hereafter called Stram) comprises 240 nuclear-encoded proteins from 112 stramenopile taxa, resulting in a concatenated alignment of 103,167 amino acid (AA) sites. The second dataset (hereafter called Ochro) removed long-branching and early-diverging heterotrophic lineages and retained 94 taxa from the more encompassing Stram alignment, including all ochrophytes, *Actinophrys sol*, Pseudofungi, Pirsoniales, and Developea. The 240 sets of orthologs were realigned, resulting in a concatenated alignment of 102,590 AA sites. Tree reconstruction for both datasets was performed with the maximum likelihood (ML) method under the site-heterogeneous LG+G4+C60 model (Wang et al. 2018), with 500 non-parametric PMSF bootstrap replicates, producing a well-supported phylogeny with all nodes having high bootstrap values (>88BS Ochro; > 94BS Stram) (fig. 1; fig. S2). Bayesian inference (BI) was conducted on both datasets using PhyloBayes-MPI v1.5 (Lartillot et al. 2013) under the CAT+GTR+G4 model. We note that the four independent chains did not converge after >12,000 iterations for either of the two datasets, but the resulting trees (table S2) are topologically highly congruent with the ML trees in both datasets. There were some minor differences in relationships among branching order among the members making up the Pelagophyceae and the Pinguiophyceae terminal taxa in the Stram datasets. Additionally, one chain of the Ochro dataset differed from the respective ML tree (fig. 1) in the position of *A. sol* and Eustigmatophyceae, recovering them as monophyletic and branching sister to SII (table S2).

**Fig. 1.**
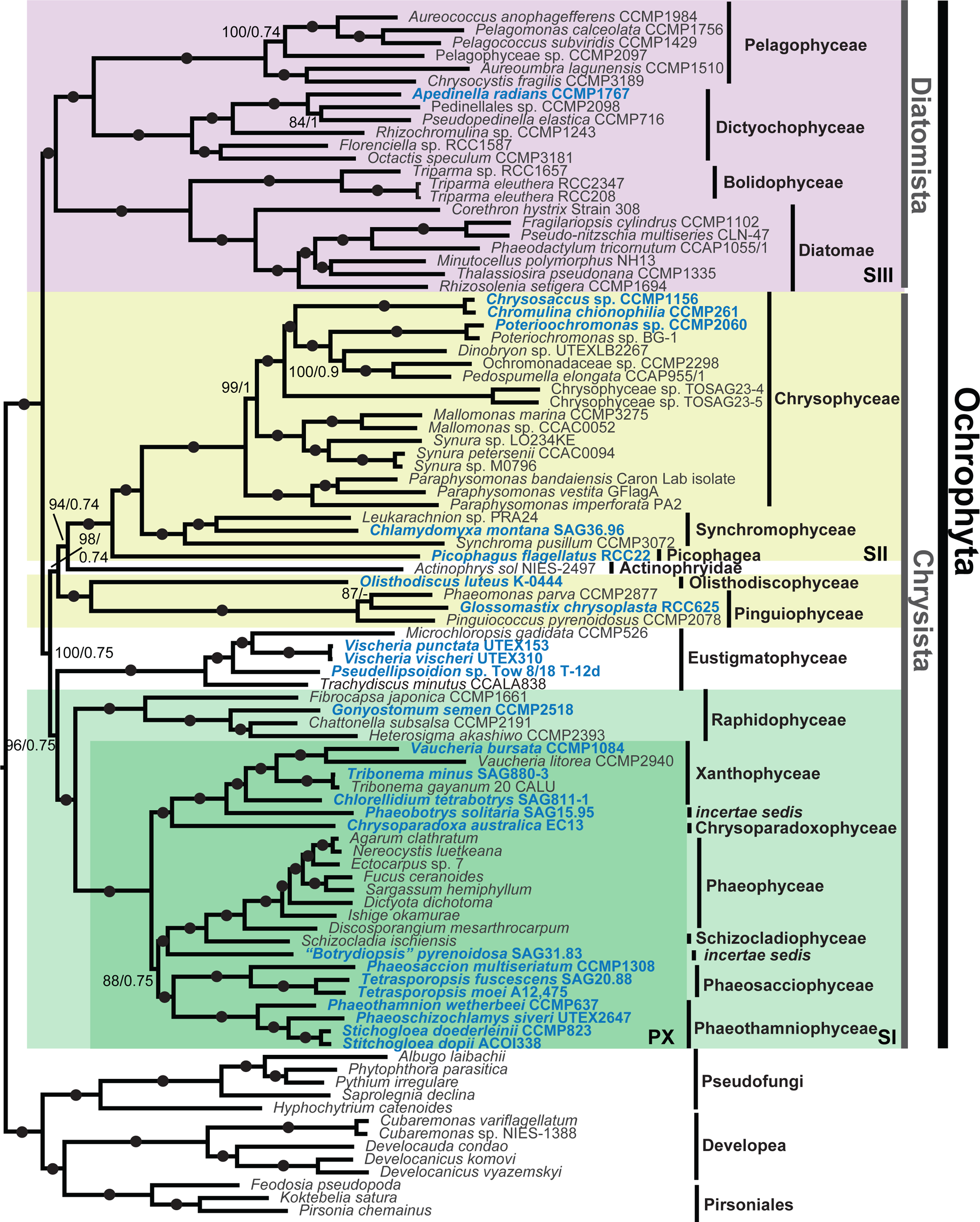
Maximum likelihood phylogeny of ochrophytes recovered from 94 taxa, 240 nuclear-encoded proteins, and 102,590 sites (Ochro dataset). Pseudofungi, Developea, and Pirsoniales were used as outgroup taxa. The previously recovered SI, SII and SII clades are resolved, with the plastid-lacking taxa *A. sol* and *P. flagellatus* notably nested within the SII clade. Quotations around *Botrydiopsis*, indicate provisional status of this taxon. Non-parametric PMSF bootstrap support (BS) values (n=500) and PhyloBayes posterior probabilities (PP) are shown on the branches as follows: BS/PP. Branches with black circles received maximum support in both analyses. Ochrophytes newly sequenced in this study are bolded and colored blue.

The relationships among stramenopile lineages, inferred from the Stram dataset (fig. S2), were highly congruent with the results of previous studies (e.g. Azuma et al. 2022; Cho et al. 2022) including the resolution of Ochrophyta as a sister group to a clade combining Pseudofungi, Developea, and Pirsoniales. All ochrophyte classes that are comprised of more than a single taxon in our dataset, including those missing in previous phylogenomic studies, such as the Phaeothamniophyceae and the Phaeosacciophyceae, were recovered as monophyletic with full support in both ML and BI methods and with both datasets (fig. 1; fig. S2). Additionally, we confidently placed three monotypic classes, Picophagea (*Picophagus flagellatus*), Olisthodiscophyceae (*Olisthodiscus luteus*), and Schizocladiophyceae (*Schizocladia ischiensis*), within the larger framework of the ochrophytes. The relationships among the ochrophyte classes were generally in agreement with previous phylogenies, including the split of Ochrophyta into two principal clades labeled Diatomista (equal to the SIII clade by Yang et al. 2012) and Chrysista (combining the SI and SII clades of Yang et al.) by Derelle et al. (2016).

Notably, some relationships that were previously weakly or moderately supported by smaller phylogenetic datasets are now robustly resolved in our analyses. For example, we recovered Olisthodiscophyceae with full statistical support as a lineage sister to Pinguiophyceae. This relationship has previously only been seen with low to moderate support in trees based on datasets of concatenated plastid genome-encoded proteins (Barcytė et al. 2021; Barcytė et al. 2022). Likewise, we have full support for the coccoid alga “*Botrydiopsis*” *pyrenoidosa* to represent an independent class-level lineage, sister to Phaeophyceae and Schizocladiophyceae, instead of being nested in the class Xanthophyceae (where it is formally classified together with genuine members of the genus *Botrydiopsis*). This is consistent with a previous three-gene analysis, which recovered a similar topology with lower statistical support (Maistro et al. 2009). Our analysis additionally clarified the position of the recently described class Phaeosacciophyceae. Whereas a previous five-gene analysis suggested that this class might be related to Xanthophyceae and Chrysoparadoxophyceae (Graf et al. 2020), our trees show the class to be robustly affiliated with Phaeothamniophyceae. Furthermore, our analyses provided full statistical support for the existence of a clade that we here treat as the class Synchromophyceae, expanded beyond its current scope.

### Topology testing reveals a rogue ochrophyte lineage

Two aspects of our trees call for further scrutiny, as they represent an incongruence with some previous results and/or point to a conflicting signal in the data. Both involve the relationships of taxa previously classified as part of the SII clade by multigene phylogenies dominated by, or exclusively relying on, plastid genome-derived sequences (Yang et al. 2012; Barcytė et al. 2021; Barcytė et al. 2022; Di Franco et al. 2022). Considering those previous analyses, Eustigmatophyceae are expected to form an internal branch of the SII clade, specifically affiliated to a subclade containing the Chrysophyceae (incl. Synurophyceae), Synchromophyceae, and potentially *P. flagellatus* (which is absent in plastid genome-based phylogenies due to its lack of a plastid genome). Both the ML and BI analyses of the Ochro dataset, and the ML analysis of the Stram dataset, placed Eustigmatophyceae as a sister lineage of the SI clade, albeit lacking full statistical support (Ochro: 94 BS/0.74 PP, Stram: 96 BS). The BI consensus of the Stram dataset placed Eustigmatophyceae as a sister lineage of a clade uniting all other SII taxa (i.e., including Pinguiophyceae and Olistodiscophyceae) plus *A. sol* without support (0.5 PP) (table S2). Interestingly, in the BI analyses of both the Ochro and the Stram dataset, one of the four chains recovers a clade containing *A. sol* and eustigmatophytes. This relationship remains a possibility, however this also may be an artifact of long branch attraction (LBA).

The second incongruence with previous results is the placement of *A. sol*, which Azuma et al. (2022) recovered as diverging prior to the ochrophyte radiation. In contrast, our trees resolve this organism as an internal branch of ochrophytes, placing it within Chrysista, and specifically within the SII clade, sister to a subclade comprised of Chrysophyceae, Synchromophyceae, and *P. flagellatus*. However, this position of *A. sol* is not fully supported in any of the analyses (94/0.74 in both Ochro and Stram datasets; fig. 1 and fig. S2). Whereas the monophyly of Chrysista, including *A. sol,* is fully supported only in the ML analyses, the BI analyses yielded support values of 0.75 and 0.87 and in the Ochro and Stram datasets, respectively. This alternative position of *A. sol* as recovered by our analyses has important consequences for the interpretation of the plastid evolution in the actinophryid lineage and is addressed in further detail below.

Given the incomplete resolution of our phylogenies and the conflicts with some of the previously reported results, we further investigated the support and robustness of our topologies by subjecting both our datasets to secondary analyses. To test how systematic errors such as LBA influenced these topologies, the secondary analyses including serial removal of fast evolving sites (FSR) and heterotacheous sites (HSR) were performed on both datasets (fig. 2). The 3,000 fastest, or most heterotacheous, sites were removed at each step and maximum likelihood trees were inferred from the resulting series of alternative datasets. These trees were reconstructed under the LG+G+C20 model with 100 nonparametric PMSF bootstraps based on these subsets.

**Fig. 2.**
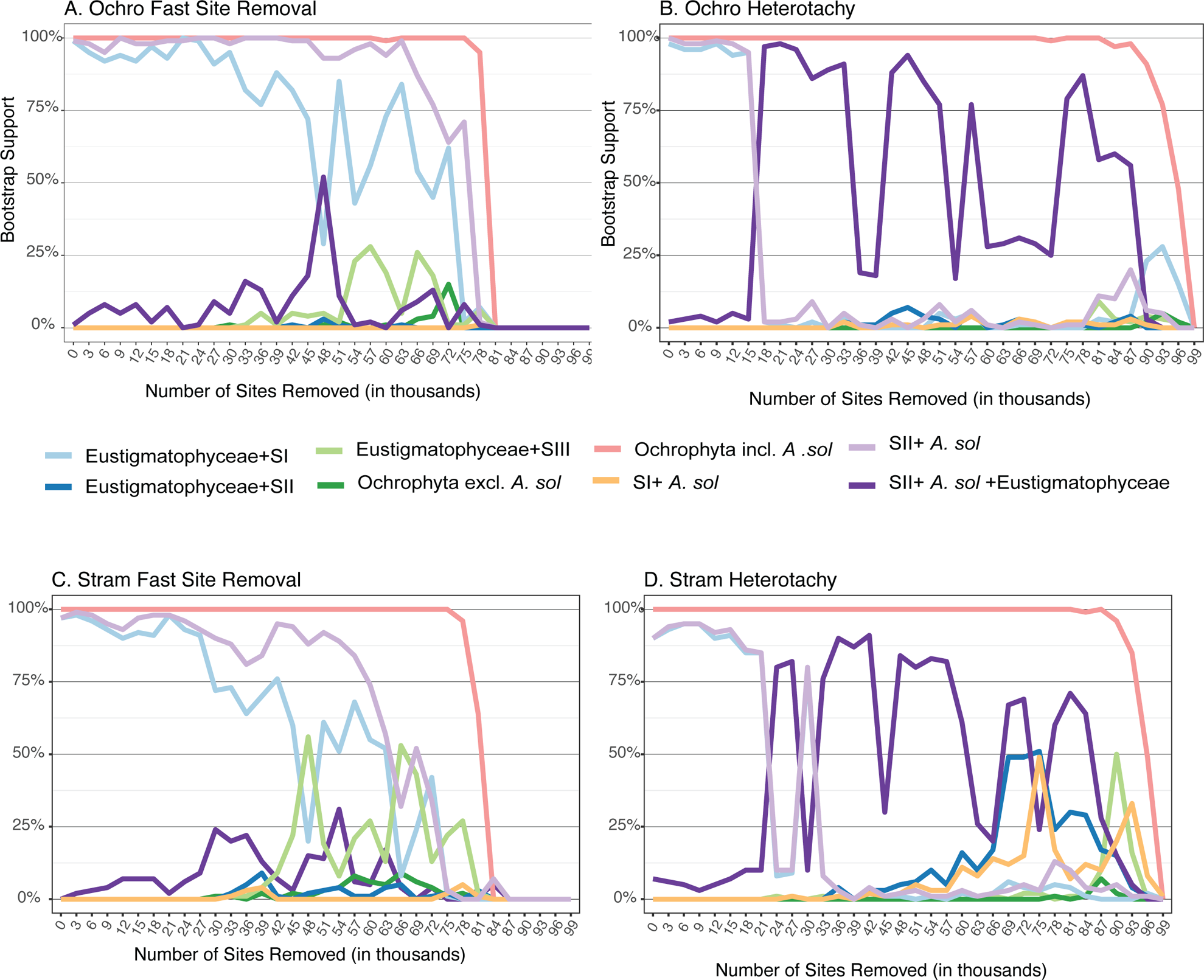
Bootstrap support for key nodes of the ochrophyte phylogeny in response to the serial removal of the 3,000 fastest evolving sites and 3,000 most heterotacheous sites until all sites were removed from each dataset. In both the Ochro dataset (A-B) and the Stram dataset (C-D), support was consistently highest for the inclusion of *A. sol* within the ochrophytes. Fast site removal generally favored a topology where SII branched with *A. sol* and SI was sister to Eustigmatophyceae. Removal of heterotacheous sites generally favored a topology uniting *A. sol*, SII and Eustigmatophyceae. Number of sites removed in thousands are on the x-axis, while non-parametric bootstrap support (n=100) are shown on the y-axis.

For the Ochro dataset we recovered full support for monophyly of ochrophytes plus *A. sol* in both the FSR and HSR analyses (fig. 2A, B). While the trees were generally stable to site removal, there were nodes with variable support, which seemed to be primarily driven by movement of eustigmatophytes as a deeply branching ochrophyte lineage. In the FSR datasets, Eustigmatophyceae was generally branching with SI, except for the dataset with 48,000 sites were removed, when support favored *A. sol,* eustigmatophytes, and SII branching together (fig. 2A). This was the only dataset where eustigmatophytes were recovered in SII, otherwise eustigmatophytes were supported as a clade with SI. Also, in this analysis, the support for *A. sol* branching with, and mostly within, SII was strong until the removal of 81,000 sites, when support for all relationships began to disintegrate (fig. 2A). In the HSR analysis, after 18,000 sites were removed, there was a shift in the placement of eustigmatophytes from branching with SI to branching with SII and *A. sol* (fig. 2B). Subsequent removals showed variable support for the placement of eustigmatophytes, rendering *A. sol* plus SII non-monophyletic, and even branching within SII at the removal of 30,000 sites (fig. 2B). With the removal of 36,000-39,000 sites, two classes that are generally a part of SII (Pinguiophyceae and Olisthodiscophyceae) moved to branching sister to SI, which drove a major loss of support for SII plus *A. sol* plus Eustigmatophyceae (fig. 2B). These conflicting topologies continued to bounce back and forth between each other throughout the rest of the HSR datasets, highlighting the uncertainty in the branching order in the early phylogeny of Chrysista.

Applying the FSR and HSR analyses to the Stram dataset yielded results generally consistent with the respective analyses of the Ochro dataset (fig. 2C, D). The monophyly of ochrophytes excluding *A. sol* was never supported in these analyses, indicating that the topology we recovered in our ML/BI analyses (fig. 1; fig. S2) is extremely robust to the removal of sites most likely to contribute to an artifactual topology. Also like in the Ochro dataset, the placement of eustigmatophytes in the Stram dataset seemed to be the driving force behind variable support for relationships among ochrophyte lineages. For example, in the FSR analysis, the position of eustigmatophytes switched from having an affinity to SI, to alternating between SI and SIII after the removal of 45,000 sites (fig. 2C). In the HSR analysis, eustigmatophytes started alternating between branching with SI, or with SII plus *A. sol*, with the removal of only 24,000 sites. After the removal of 33,000 sites, a clade containing Eustigmatophyceae, *A. sol*, and SII was the most supported topology for the eustigmatophytes (fig. 2D). In the FSR analysis, *A. sol* remained a well-supported member of SII until more than 60,000 sites were removed (fig. 2C). In both analyses there was little to no support for Eustigmatophyceae branching with SII (to the exclusion of *A. sol*), highlighting the instability of Eustigmatophyceae driving variable support for the conflicting topologies.

As an independent way to assess how our sequence datasets supported the inferred topologies, we subjected them to random gene resampling (RGS) analysis (fig. S3). For both datasets independently, genes were randomly sampled at intervals of 20%, 40%, 60% and 80% of the complete dataset until meeting the 95% confidence interval that all genes were included in a subsampled dataset (20% n=14; 40% n=6; 60% n=4; 80% n=2). Trees were inferred for each alternative subsample, and statistical support for relationships of special interest were summarized across the subsamples. For both datasets, the highest degree of variability was seen in the smallest dataset (20% of total genes) and the placement of Eustigmatophyceae remained unclear in all subsampled analyses. In the Ochro dataset, monophyly of SII, *A. sol* and Eustigmatophyceae, as well as Eustigmatophyceae plus SI, both received highly variable support in the datasets comprised of 20% of the genes (fig. S3A). As the number of genes increased, so did the support for SII plus *A. sol* plus Eustigmatophyceae, yet even with 80% of the genes included in the datasets, support wavered, leaving the placement of the eustigmatophytes unresolved. In the Stram dataset, Eustigmatophyceae were variably supported as monophyletic with SI in the 20% dataset, and support for this relationship strengthened as more genes were included (fig. S3B).

As an alternative to the gene concatenation-based approaches, we additionally analyzed both datasets using the coalescence-based phylogenomic analysis implemented in ASTRAL-III (Zhang et al. 2018). The backbone of the ochrophytes phylogeny was poorly supported in this analysis but topologically consistent with the results of concatenation-based analyses. *Actinophrys sol* was either sister to or nested within the SII clade in both cases. Although the exact placement of *A. sol* within SII was not well supported in these trees (Ochro: 0.56; Stram: 0.7), there was full support for *A. sol* being a member of Ochrophyta (table S2). The position of Eustigmatophyceae also differed between the two datasets, with this lineage being sister to the SI clade (Ochro dataset, support value of 0.41) or sister to SII plus *A. sol* (Stram dataset, support value of 0.45).

Finally, we used the approximately unbiased (AU) test (Shimodaira 2002) to evaluate our Ochro ML topology (fig. 1) and nine alternative topologies (fig. 3A). The alternative topologies to the ML topology (fig. 3B) included *A. sol* branching sister to ochrophytes (fig. 3C), monophyly of *A. sol* and the SI clade (fig. 3D) as well as seven other topologies specifically regarding the placement of either *A. sol* or Eustigmatophyceae (fig. 3E-K). Four of the ten topologies were not rejected, all differing in the position of eustigmatophytes. These included the recovered ML topology, which has eustigmatophytes branching sister to SI (fig. 3B), and three topologies where *A. sol*, Eustigmatophyceae, and SII taxa are together monophyletic but in different branching orders (fig. 3G-I).

**Fig 3.**
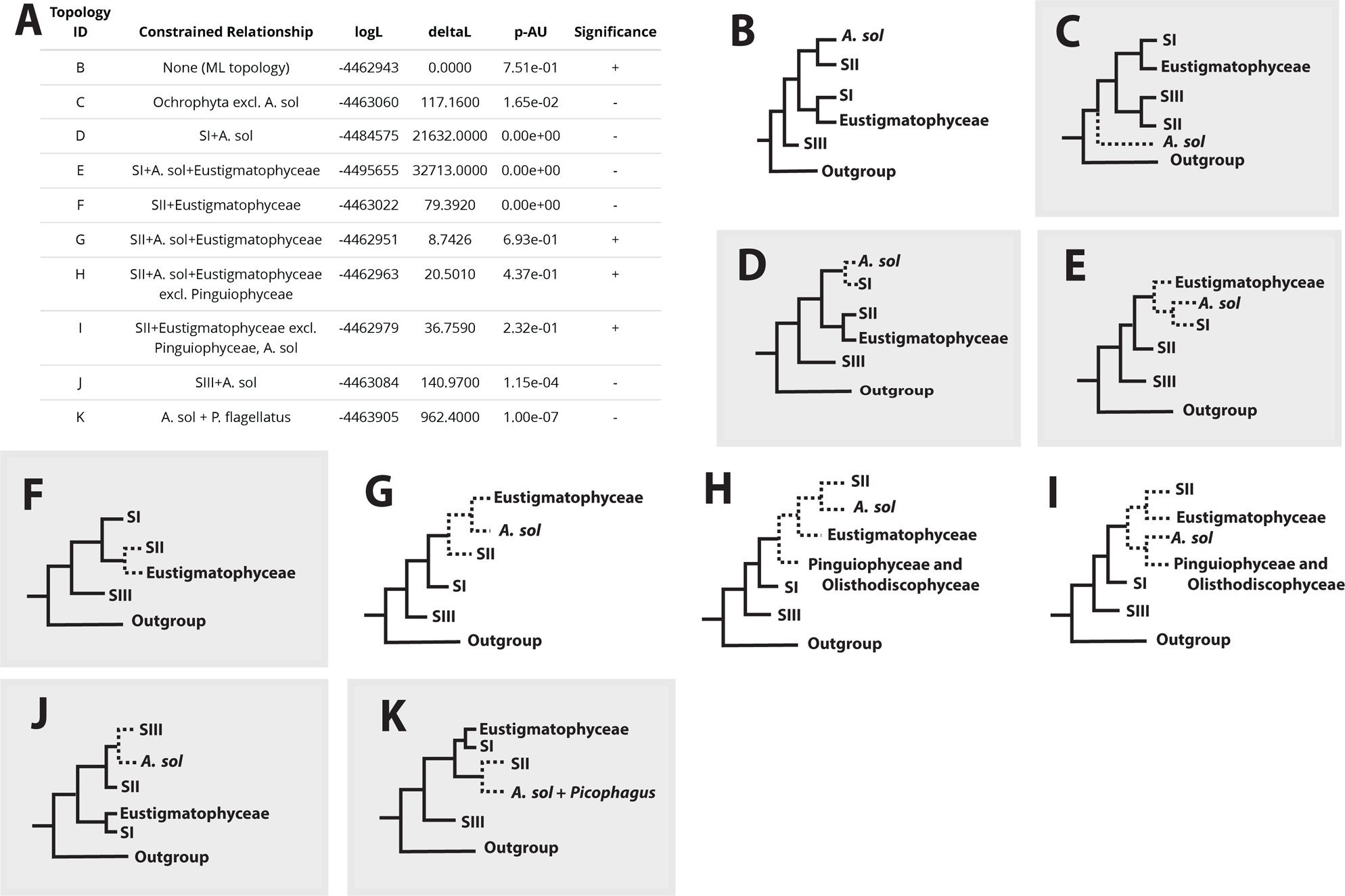
Approximately unbiased test results for possible ochrophyte topologies. A) The list of constraints placed on 10 different potential topologies, with the ‘+’ denoting relationships that were not rejected and a ‘-‘ indicating topologies that were rejected. B) Schematic of the unconstrained tree recovered by our analysis (i.e. that in fig. 1). C-K) Schematics of each recovered alternative topology that was tested, where dotted lines indicate the constrained nodes and solid lines represent unconstrained nodes.

Thus, despite the major improvement of the taxon sampling achieved by this study, this result further highlights the instability of the position of Eustigmatophyceae among ochrophytes and they may be considered a rogue class (Sanderson and Shaffer 2002). Topologies recovered in previous studies are conflicting, depending upon whether the genes used were located on either the plastid or the nuclear genome. When plastid genes are used, there is overwhelming evidence for eustigmatophytes being nested within the SII clade, specifically related to Chrysophyceae and Synchromophyceae (Yang et al. 2012; Ševčíková et al. 2016; Barcytė et al. 2022; Barcytė et al. 2024). Analyses of nuclear genes tend to favor eustigmatophytes being a sister lineage of the SI clade (this study; Noguchi et al. 2016; Leonard et al. 2018; Dorrell et al. 2021; Cho et al. 2022), However, with these test results, we are unable to reject the topology recovered by the plastid genes (fig. 3H-I). This conflict was previously also observed and analyzed by Di Franco et al. (2022), who concluded that the plastid genome-based phylogenies are more credible. Strikingly, their mitochondrial genome-based phylogeny, although sparsely sampled and poorly supported, recovered eustigmatophytes being the SI sister lineage, like the nuclear genes-based phylogeny. Thus, we refrain from making any conclusions regarding the position of eustigmatophytes in the ochrophyte phylogenetic tree and call for further investigations into the reasons for the peculiar behavior of this lineage in phylogenetic analyses.

### Actinophrys sol represents a complete loss of the ochrophyte plastid

Ochrophyta are generally accepted to have acquired a red algal-derived plastid through a higher-order endosymbiosis, although the exact origin of the ochrophyte plastid has been contested for decades (Dorrell and Bowler 2017). The chromalveolate hypothesis posited that the ochrophyte plastid evolved vertically from a plastid-bearing ancestor shared by several other eukaryote groups with complex plastids of rhodophyte origin, including cryptophytes, haptophytes, and myzozoans (Cavalier-Smith 1999). This scenario would necessitate that all stramenopile lineages would have emerged from plastid-bearing ancestors. However, genomic studies and systematic investigations of heterotrophic stramenopiles, including those from lineages phylogenetically closest to ochrophytes, favor a hypothesis of a more recent plastid gain within stramenopiles, which gave rise to ochrophytes (Stiller et al. 2014; Dorrell et al. 2017; Strassert et al. 2021). Within this context, the phylogenomic placement of *Actinophrys sol* (representing a broader group, Actinophryidae) as a lineage sister to ochrophytes is notable (Azuma et al. 2022). Analyses of *A. sol* genomic and transcriptomic data presented by the authors strongly supported the absence of a plastid in this organism, yet revealed a dozen organellar aminoacyl-tRNA synthases of putative algal origin, suggesting the presence of a photosynthetic endosymbiont in its ancestor. Although admitting the possibility of a *bona fide* plastid loss in the *A. sol* lineage, Azuma et al. (2022) favored a scenario in which the plastid had not been stably integrated into the host cell prior to the divergence of Actinophryidae and Ochrophyta, but the common ancestor of both groups already possessed nuclear genes for plastid functions acquired by horizontal gene transfer (HGT) from algal sources.

The feasibility of such a scenario of the plastid history in actinophryids has been supported by a recent study documenting the acquisition of nuclear genes for plastid-targeted proteins preceding the establishment of the Pyramimonadales-derived plastid in euglenophytes (Karnkowska et al. 2023). However, the monophyly of ochrophytes, to the exclusion of *A. sol,* received strong support only in the concatenation-based analysis reported by Azuma et al. (2022), whereas their coalescence-based analysis showed a much lower support for it (Azuma et al. 2022). Our analyses do not reproduce those results, as both ML and BI analyses of concatenated sequence datasets as well as the coalescence-based approach all place *A. sol* into ochrophytes, consistently into the Chrysista clade and in most analyses (except for the ASTRAL analysis of the Stram dataset) as a sister lineage of a subclade including Chrysophyceae, Synchromophyceae and *P. flagellatus*.

To test our hypothesis that the discrepancy between the results and those of Azuma et al. (2022) stem from the increased taxon sampling, we constructed one additional dataset, from here forward called Act. This dataset is comprised of the same 75 taxa from Azuma et al. (2022), however with our ortholog selection that was completed using increased taxonomic breadth, which may have enabled the identification of previously undetectable mid-paralogs. The ML analysis of this 95,605 AA site dataset recovered *A. sol* sister to ochrophytes as in Azuma et al., with the ochrophyte monophyly (to the exclusion of *A. sol*) highly supported (BS 94; fig. S4). The BI analysis of this dataset (which was not done by Azuma et al.) yielded *A. sol* and three separate ochrophyte lineages in an unresolved polytomy (table S2). The ASTRAL analysis of the Act dataset directly conflicted with the result of Azuma et al., placing *A. sol* (albeit with poor support) among ochrophytes, in a position equivalent to the one recovered in most of our analyses (table S2). It thus seems that the substantially improved taxon sampling in our datasets, including representatives of deeply diverged ochrophyte lineages (*O. luteus*, *P. flagellatus*), as well as outgroup taxa (Pirsoniales), helps in extracting phylogenetic signal from the data and stabilizes the results.

Despite variability in some characteristics, such as pigment composition and metabolic pathways (Dorrell et al. 2017; Nonoyama et al. 2019), plastids across all ochrophyte lineages are undoubtedly from a single origin. In addition to the strong support for the monophyly of ochrophyte plastids provided by plastid genome-based analyses (e.g., Muñoz-Gómez et al. 2017), the ochrophyte plastids retain highly conserved ultrastructural and molecular features, including membrane organization and the mechanism of protein import (Dorrell and Bowler 2017). Based on our phylogenomic analysis, we reinterpret *A. sol* as a *bona fide* ochrophyte lineage that evolved from an ancestor possessing a fully integrated complex plastid, and hence a case of complete plastid loss in ochrophytes. Additional genomic data will be required from *A. sol* relatives, including *Actinosphaerium* species, to test the robustness of this conclusion. Furthermore, those data will be instrumental to potentially break the long, deeply-diverging, branch of *A. sol*, and may further stabilize the phylogenetic position of actinophryids. Nevertheless, *A. sol* demonstrates that plastid loss is evolutionarily feasible in Ochrophyta. Strikingly, a second ochrophyte, newly sampled here, may share the same novel characteristic.

### *Picophagus flagellatus* represents another plastid loss within ochrophytes

*Picophagus flagellatus* is a marine heterotrophic nanoflagellate that was originally classified within the Chrysophyceae based on an 18S rDNA tree (Guillou et al. 1999). Later, Cavalier-Smith and Chao (2006) produced an 18S rDNA tree that recovered a weakly supported clade containing *P. flagellatus* and *Chlamydomyxa labyrinthuloides*. Based on this topology they defined a new class, Picophagea, to accommodate both species. However, this poorly resolved relationship was not recovered in subsequent phylogenies based on the 18S rDNA sequences (Grant et al. 2009; Patil et al. 2009; Schmidt et al. 2015). In addition, *P. flagellatus* is not united with *C. labyrinthuloides* by any obvious morphological feature. In our expanded phylogenomic dataset we recovered *P. flagellatus* as sister to a fully supported clade comprised of Chrysophyceae and the class Synchromophyceae, here expanded to include *C. labyrinthuloides* (fig. 1; fig. S1). We adopt the class Picophagea to classify *P. flagellatus* as a higher-order taxon, but restrict its definition such that it is currently monotypic.

Investigation by transmission electron microscopy (TEM) revealed no cellular structures in *P. flagellatus* that could be recognized as candidates for a plastid (Guillou et al. 1999), however, this does not necessarily mean that the plastid is truly absent. Organisms including *Euglena longa* (Füssy et al. 2020) and the ochrophyte *Leukarachnion* sp. PRA-24 (Barcytė et al. 2024) have no clear ultrastructural evidence for a plastid, yet retain strong genomic evidence that the organelle persists. Most notably, both taxa maintain a plastid genome. To investigate whether *P. flagellatus* retains a cryptic plastid, we generated a draft genome assembly for this organism, comprised of ∼50.9 Mbp distributed in ∼21,700 scaffolds. The assembly statistics are influenced by a proportion of the scaffolds being derived from bacterial contamination, but manual inspection indicated that the *P. flagellatus* nuclear genome is represented by scaffolds with ∼11.7× read coverage and typically tens of thousands bp long. Assessing the assembly completeness with BUSCO (95% complete, using the predefined Stramenopiles lineage dataset as a reference) gives comparable values to ochrophyte genomes considered to be completely sequenced (97%-100% complete BUSCOs; fig. S5). In addition to the nuclear genome sequences, the assembly contains a high-coverage scaffold (∼223.4×; NODE_359) that represents a circular-mapping (i.e. presumably complete) mitochondrial genome sequence. Hence, the fact that we found the *P. flagellatus* genome assembly to be devoid of any sequences that would be derived from a plastid genome can be interpreted as a strong indication the plastid genome is truly absent from this organism.

This finding alone does not rule out the possibility of *P. flagellatus* possessing a genome-lacking plastid, as is found in the colorless chrysophyte genus *Paraphysomonas* (Dorrell et al. 2019) and various taxa outside ochrophytes (e.g., Smith and Lee 2014; Gawryluk et al. 2019; Janouškovec et al. 2019; Mathur et al. 2023). However, our analysis of the *P. flagellatus* transcriptome and draft genome assembly did not reveal any homologs of the components of the plastid protein complexes that mediate translocation of imported proteins across different plastidial membranes (SELMA, TOC and TIC complexes; Maier et al. 2015). Additionally, homologs for the stromal processing peptidase required for maturation of imported proteins by removing the N-terminal transit peptide (Huesgen et al. 2013) were missing from these data. Additionally, the dynamin-related protein DRP5B and plastidial (i.e. Cyanobacteria-derived) versions of FtsZ making up the plastid division machinery (Miyagishima et al. 2014) were not detected. In contrast, at least some of these proteins are readily identified in non-photosynthetic ochrophytes where the plastid is retained, such as *Leukarachnion* sp. PRA-24 and even in the highly reduced plastids of *Paraphysomonas* spp. (Dorrell et al. 2019; Barcytė et al. 2024; see also table S3). The lack of these proteins in *P. flagellatus* does not seem to result from incomplete data or inadequate searching procedure, as proteins analogously servicing the mitochondrion were identified (table S4). Therefore, these results make it highly unlikely that *P. flagellatus* harbors even the most reduced form of a plastid.

The conclusion that a plastid is completely absent in *P. flagellatus* is further supported by our analysis of enzymes of metabolic pathways typically localized in ochrophyte plastids. In contrast to plastid-bearing non-photosynthetic ochrophytes, including *Paraphysomonas* spp. with their extremely reduced plastids (Dorrell et al. 2019), no genes for the heme biosynthesis pathway could be detected in the *P. flagellatus* sequence data. Despite an inability to synthesize heme, numerous homologs of heme-utilizing proteins were detected (table S4), suggesting that *P. flagellatus* acquires heme from food. Other hallmark plastidial metabolic pathways, retained in different combinations by non-photosynthetic plastids across eukaryotes (e.g., Salomaki et al. 2015; Dorrell et al. 2019; Salomaki and Kolisko 2019; Füssy et al. 2020; Kayama et al. 2020; Pánek et al. 2022; Mathur et al. 2023; Barcytė et al. 2024), are likewise completely missing in *P. flagellatus*. These include the DOXP pathway for the biosynthesis of isoprenoid precursors, the type II fatty acid synthesis (FAS) pathway, the shikimate pathway and its downstream branches leading to aromatic amino acids or folate, enzymes for the synthesis of plastid-specific structural lipids (galactolipids, sulfoquinovosyldiacylglycerol), or pathways for the synthesis of plastidial terpenoid quinols and their derivatives (plastoquinol, phylloquinol, tocopherols/tocotrienols). Some of the respective products are essential and, like heme, are likely acquired by *P. flagellatus* from food (aromatic amino acids, folate) or synthesized by alternative non-plastidial pathways (isoprenoid precursors synthesized by the mevalonate pathway, fully conserved in *P. flagellatus*; table S4).

As an alternative to the missing plastid-associated type II FAS enzymes we identified a homolog of the large multidomain type I fatty acid synthase (FAS I) in *P. flagellatus* (table S4). Phylogenetic analysis revealed that the *P. flagellatus* FAS I protein forms a clade with homologs from related taxa, including *Leukarachnion* sp. PRA-24 and several photosynthetic and non-photosynthetic chrysophytes (fig. 4). The occurrence of FAS I in ochrophytes has not been, to our knowledge, reported before and the exact origin of this gene is unclear, but its absence in other ochrophytes suggests it was acquired by HGT by the common ancestor of *P. flagellatus* and Chrysophyceae/Synchromophyceae. The acquisition of FAS I rationalizes the absence of type II FAS pathway in *Leukarachnion* sp. PRA-24 (Barcytė et al. 2024) and colorless chrysophytes (Dorrell et al. 2019), and might be a predisposition to plastid reduction, which is particularly common in the respective ochrophyte clade compared to other lineages.

**Fig. 4.**
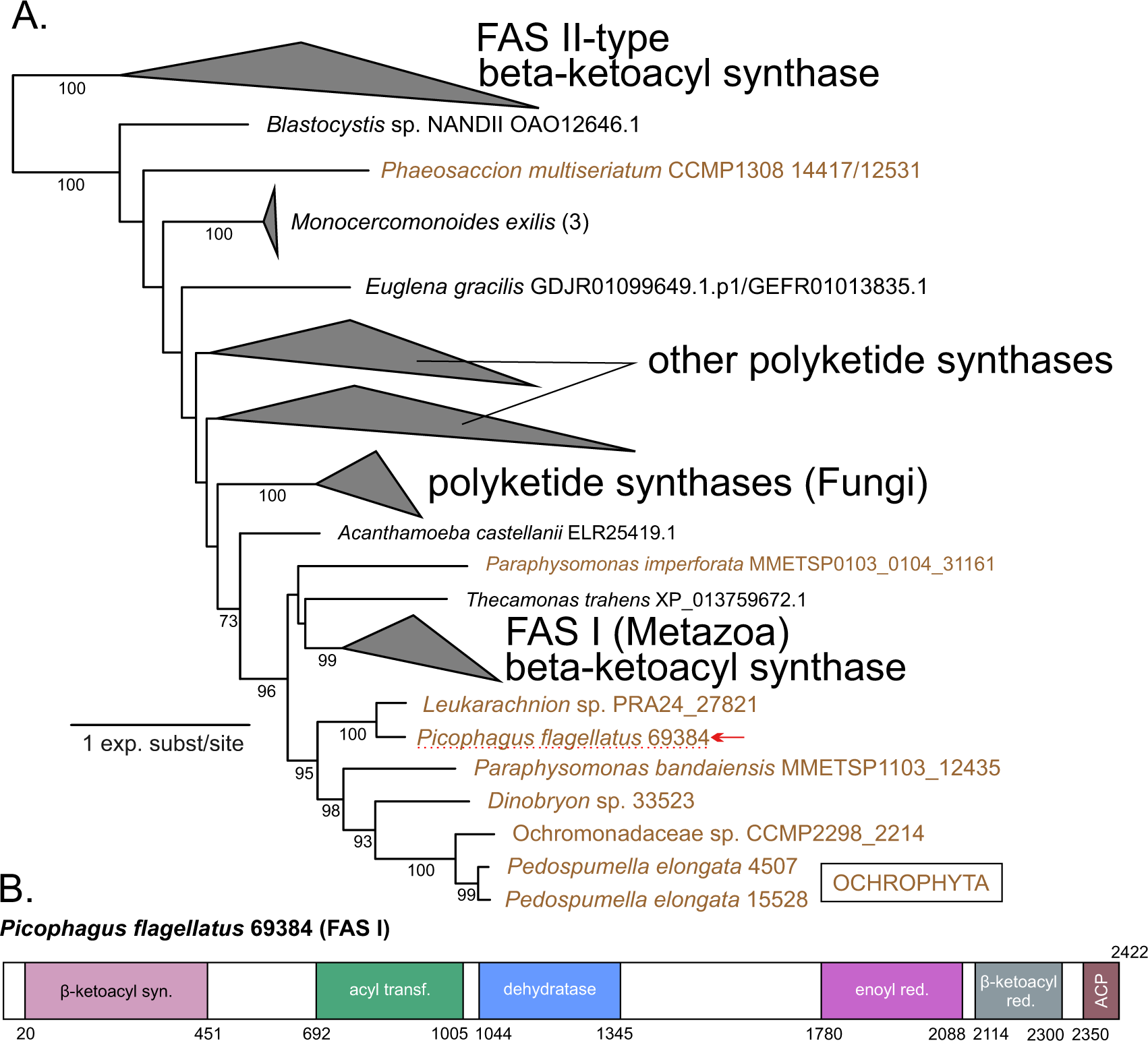
Phylogenetic position and domain architecture of a putative type I fatty acid synthase (FAS I) identified in several ochrophytes including *Picophagus flagellatus*. A) Phylogenetic analysis of FAS II-type stand-alone β-ketoacyl synthases and the corresponding domains of FAS I and polyketide synthase (PKS). Only bootstrap values ≥50 are shown. Clades out of interest were collapsed as closed triangles, the tree was rooted arbitrarily on the long branch separating FAS II sequences from FAS I/PKS. B) Domain architecture of the putative FAS I protein from *P. flagellatus* (the numbers indicate coordinates in the amino acid sequence). Multiple sequence alignment contained 361 protein sequences and 368 positions after trimming, it is available in Figshare link ####.

Combined, our bioinformatic analyses build a strong case for *P. flagellatus* being a secondarily aplastidic ochrophyte. Furthermore, our phylogeny recovered *A. sol* outside a fully supported clade containing *P. flagellatus*, arguing for two independent events of plastid loss in ochrophytes. This is noteworthy, as only a few convincing cases of a plastid loss have been reported so far, one among primary plastids (in Picozoa; Schön et al. 2021) and a few others restricted to some parasitic lineages in Apicomplexa and Dinoflagellata (Zhu et al. 2000; Maciszewski and Karnkowska 2019; Salomaki and Kolisko 2019b; Mathur et al. 2023).

It was recently argued that when a plastid is lost, it leaves no genetic trace in the respective lineage (Holt et al. 2023). In this context, it is interesting to note that *A. sol* has been shown to exhibit orthologs of seven aminoacyl-tRNA synthetases (aaRSs) that seem to have been acquired from algal sources by an ochrophyte ancestor to support translation in the newly established plastid. At the same time, the newly acquired aaRSs (thanks to dual targeting) have replaced the aaRSs that had serviced the mitochondrion in stramenopile ancestors of ochrophytes (Azuma et al. 2022). We identified homologs of six out of the seven aaRSs in *P. flagellatus* and reconstructed phylogenies of them, including homologs from the new ochrophyte sequence data generated by us and from aplastidic ochrophyte relatives (Pirsoniales and Developea) reported by Cho et al. (2022). Our analyses are consistent with the previous inference that the original mitochondrial aaRSs have been replaced by the dually localized aaRSs in the common ancestor of ochrophytes and additionally showed that the respective orthologs occur also in *P. flagellatus*. This is most convincingly demonstrated (in terms of the statistical support of tree topologies) for valyl-tRNA synthetase (fig. 5) and aspartyl-tRNA synthetase (fig. S6; datasets and trees for all investigated aaRSs are available on Figshare link). Hence, the aaRSs can be viewed as a genetic legacy of the plastid in both *A. sol* and *P. flagellatus*, which is retained even when the plastid is lost, as the proteins are also essential for the mitochondrion.

**Fig. 5.**
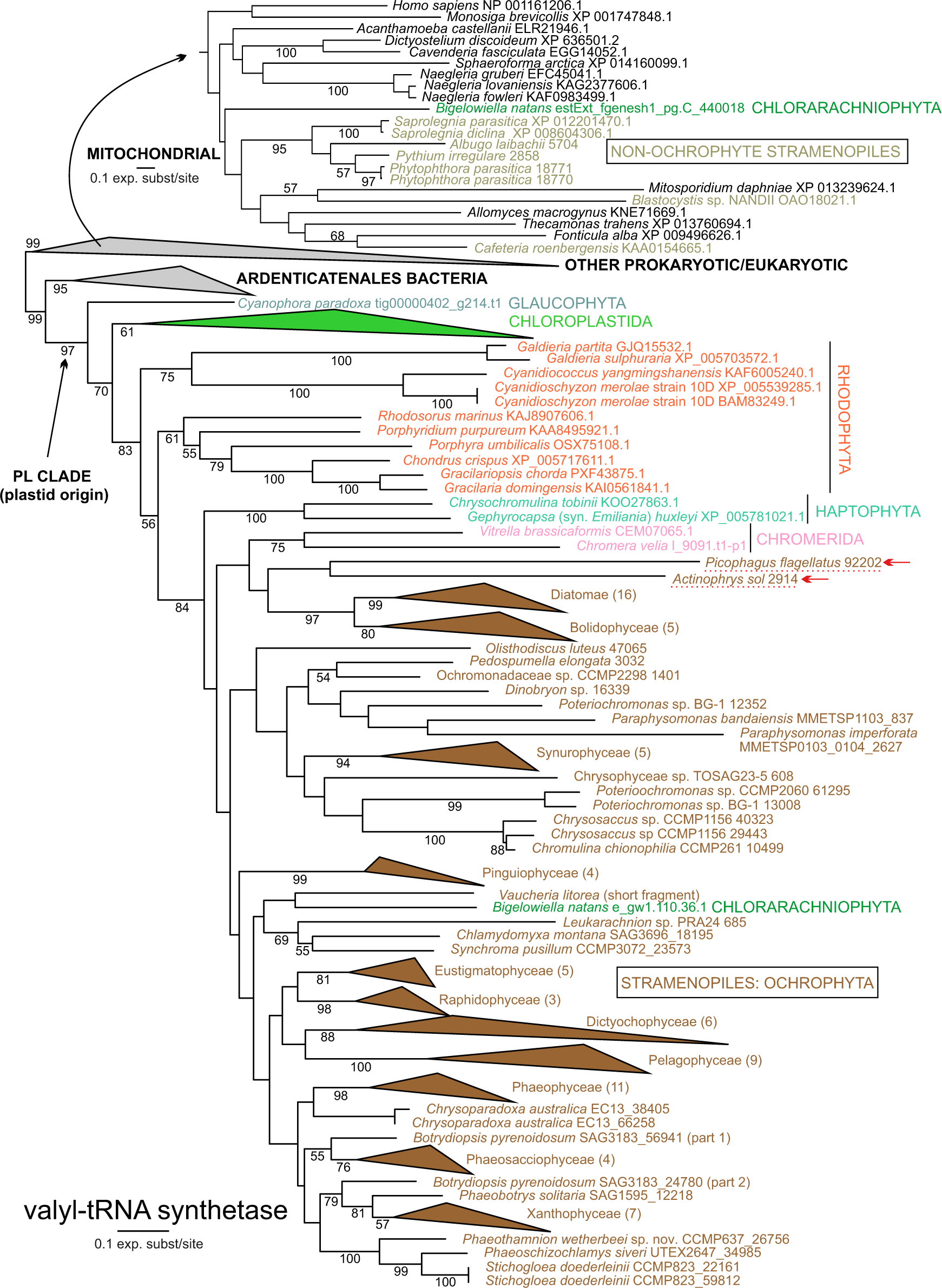
Phylogenetic analysis of organellar and cytoplasmic valyl-tRNA synthetases (ValRS) showing the position of proteins from *Picophagus flagellatus* and *Actinophrys sol* (highlighted with arrows) branching within a clade of dually-targeted enzymes of a plastid origin (unrelated to the original mitochondrion-targeted ValRS present in non-ochrophyte stramenopiles; the subtree on the top). Only bootstrap values ≥50 are shown. Clades out of interest were collapsed as closed triangles, the tree was rooted arbitrarily. The clade of mitochondrial homologs that branched within the collapsed clade is shown in detail separately. Multiple sequence alignment contained 739 protein sequences and 762 positions after trimming, it is available in Figshare link ####.

Strikingly, *P. flagellatus* also exhibits a putative remnant of an exclusively plastidial pathway. Similar to other non-photosynthetic ochrophytes (Dorrell et al. 2019; Barcytė et al. 2024), *P. flagellatus* lacks the lysine biosynthesis pathways, which in photosynthetic ochrophytes is wholly localized in the plastid (Dorrell et al. 2017). However, homologs of the final two enzymes of the pathway, diaminopimelate epimerase (DapF) and diaminopimelate decarboxylase (DAPDC), do exist in *P. flagellatus* (table S4). The former is extremely divergent (leaving its origin and functionality uncertain), and it is likely cytosolic according to the results of multiple targeting prediction programs. For the second enzyme, phylogenetic analyses are compatible with the notion that DAPDC has been retained from the photosynthetic ancestors of *P. flagellatus* (fig. S7, S8). The enzyme was predicted to be targeted to the residual plastid in the chrysophytes (Dorrell et al. 2019). The *P. flagellatus* DAPDC also exhibits an N-terminal extension compared to bacterial homologs, but it is predicted as a mitochondrion-targeting presequence by the majority of the prediction tools employed (table S4). The retention of the enzyme likely enables the organisms to utilize diaminopimelate (a precursor of both lysine and peptidoglycan plentifully produced by bacteria) for lysine production. A systematic search for genetic traces of the plastid in *P. flagellatus* may uncover additional cases, but the examples of DAPDC and aaRSs argue that the hypothesis by Holt et al. (2023) on plastids leaving no traces upon their loss may be an inappropriate generalization of the specific situation in secondarily aplastidic myzozoans.

### Redefining the class Synchromophyceae

In addition to providing insights into the plastid losses within ochrophytes, our analyses have implications for refinements to the taxonomy and classification of this group, such as preferring the historical categorization of synurophytes as a subgroup of Chrysophyceae, rather than a separate class. Here we concentrate on another case, namely the definition of the class Synchromophyceae. The class was established in 2007, to include a newly described marine amoeboid alga, *Synchroma grande* (Horn et al. 2007). In addition to its isolated phylogenetic position in ochrophytes, it exhibited an unprecedented morphological feature, termed chloroplast complexes, that were proposed as the defining characteristic of the new class. The chloroplast complexes exist where the two innermost membranes of the conventional secondary ochrophyte plastid delimit several separate compartments located in a common periplastidial compartment, thus being collectively enveloped by two membranes corresponding to the two outermost membranes. The same plastid configuration was later confirmed in another *Synchroma* species, *S. pusillum* (Schmidt et al. 2015). However, several additional organisms have emerged over the years, including *Chlamydomyxa labyrinthuloides* and *Leukarachnion* sp. PRA-24, as related to *Synchroma* in analyses of limited molecular data, yet they lack this characteristic morphology (Grant et al. 2009; Patil et al. 2009; Schmidt et al. 2015). Of these taxa, *S. pusillum* has been the only species included in phylogenomic datasets to date (Derelle et al. 2016; Tice et al. 2021; Cho et al. 2022).

To test the evolutionary relationships of *Synchroma* and other poorly studied ochrophytes, we generated the first molecular data for *Chlamydomyxa montana* SAG 36.96, which was confirmed to be a close relative of *C. labyrithuloides* by an 18S rDNA phylogeny (fig. 6; fig. S1). In addition, we included transcriptome data generated as part of a parallel study (Barcytė et al. 2024) from *Leukarachnion* sp. PRA-24, the only culture representative available of a broader ochrophyte clade that has been primarily documented by environmental DNA. This clade has shown to be specifically affiliated to a clade containing *C. montana* and *S. pusillum* in the 18S rRNA gene phylogeny (fig. 6; fig. S1). Our results corroborated, with full statistical support, the existence of a clade containing all three species, denoted SCL, and resolved their previously unclear branching order with *S. pusillum* sister to a clade comprising *C. montana* and *Leukarachnion* sp. PRA-24 (fig. 1; fig. S2). The SCL clade was in turn resolved with full support as the sister group of Chrysophyceae (including Synurales), yet deeply diverged from it (fig. 1; fig. S2).

**Fig. 6.**
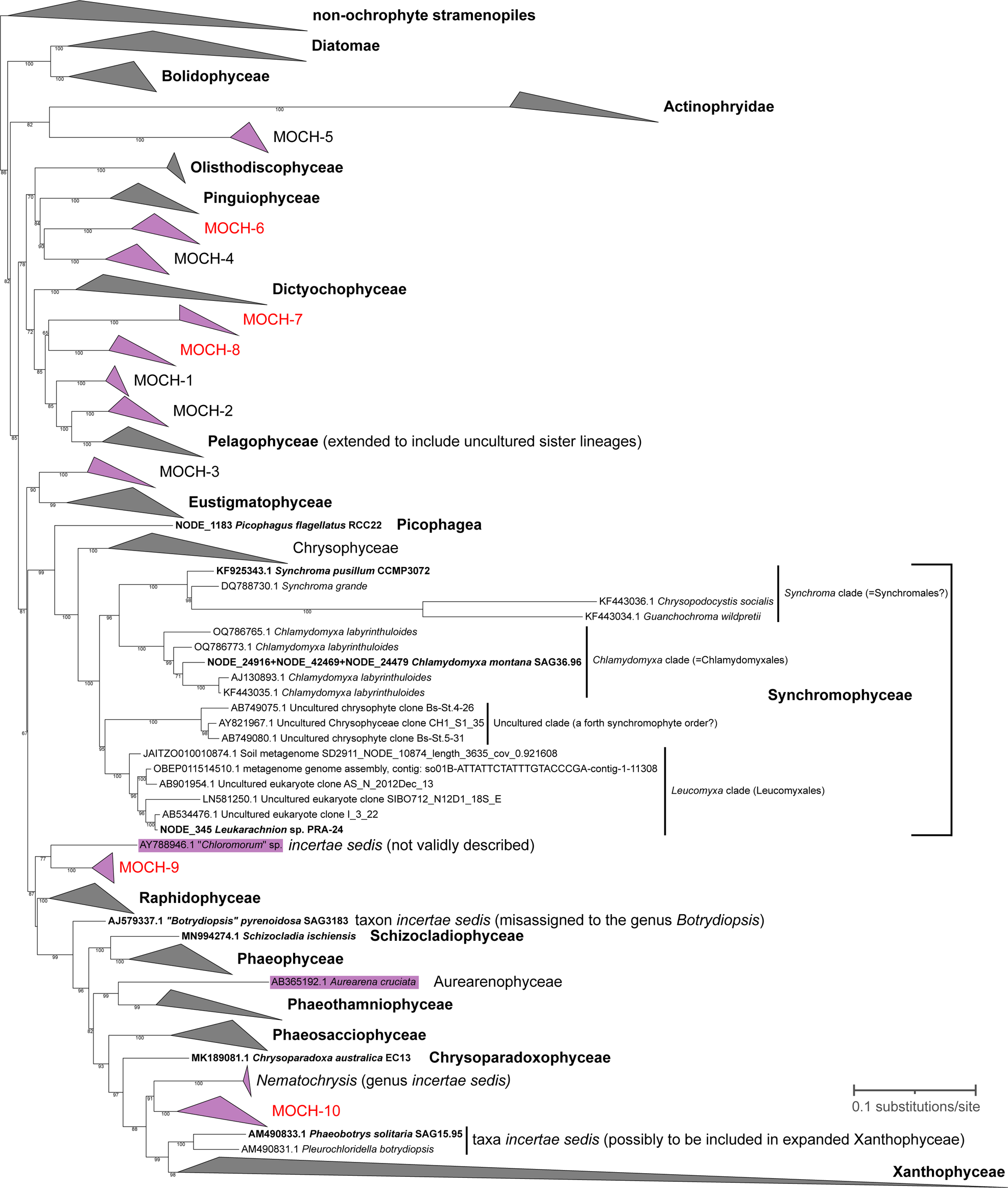
Phylogenetic tree of the 18S rDNA that identifies a dozen ochrophyte class-level lineages that are yet to be included in phylogenomic analyses. The figure was modified from the ML tree provided in full as fig. S1. The non-ochrophyte outgroup taxa and the clades representing formally recognized ochrophyte classes, except for Synchromophyceae and any monotypic classes, were collapsed and are represented by triangles. Several *incertae sedis* taxa branch outside these classes and may thus represent classes of their own. The four major clades in the class Synchromophyceae (as redefined here), including one represented only by environmental sequences from uncultured organisms, are annotated. Taxa (individual species or larger clades) represented in our phylogenomic analyses are typed in bold, whereas the major separate lineages missing from it are highlighted in purple. The latter include the monotypic class Aurearenophyceae and two of the *incertae sedis* taxa: *Nematochrysis*, proposed to be affiliated to Chrysoparadoxophyceae (Graf et al. 2020) but more closely related to the “MOCH-10” clade (see below) and Xanthophyceae in our tree; and a lineage represented in GenBank by a series of highly similar sequences attributed to the organism denoted “*Chloromorum* sp. *toxicum*”, not yet formally described or morphologically documented in the literature. In addition, ten class-level clades represented solely by uncultured marine ochrophytes (MOCH) and each robustly recovered as monophyletic, but deeply diverged from other classes or clades emerged from the analysis of presently available 18S rRNA gene sequences: MOCH-1 to MOCH-5 recognized previously (Massana et al. 2014) and MOCH-6 to MOCH-10 (highlighted in red) newly defined here. The MOCH-5 clade includes once cultured strains (represented by RCC862 in the tree) that were lost before they could be characterized (Marie et al. 2017). The origin of the the sequence branching in the MOCH-3 clade and assigned to “Chattonella subsalsa isolate BH65” in the respective database record is uncertain due to the lack of a corresponding publication, but the taxon name is clearly misassigned (*Chattonella subsalsa* is an unrelated raphidophyte; see also fig. S1).

The strong evidence for the monophyly of the SCL taxa contrasts with the unsettled classification of these organisms. It seems practical to consider the SCL clade a single class, however this has not previously been formally proposed. Schmidt et al. (2015) demonstrated that two amoeboid marine algae, *Chrysopodocystis socialis* and *Guanchochroma wildpretii*, are even more closely related to *Synchroma* than *C. montana* and *Leukarachnion* sp. PRA-24 (see also fig. 6 and fig. S1), yet they were reluctant to place them explicitly into any class. We thus propose the circumscription of Synchromophyceae be modified (see taxonomic appendix) to include all taxa of the SCL clade, including *Ch. socialis* and *G. wildpretii* missing from our analysis, yet with robust evidence closely related to *Synchroma*. All members of the redefined Synchromophyceae are united by an amoeboid (rhizopodial) morphology and the absence of flagella in vegetative stages, which contrasts to the predominant and certainly ancestral monadoid (flagellate) morphology of the sister class, Chrysophyceae. The unusual organization of the plastid that was originally considered a defining characteristic of the class Synchromophyceae is best interpreted as an evolutionary innovation (autapomorphy) of the narrow lineage represented by the genus *Synchroma*. Thus, like their sister group Chrysophyceae, the redefined Synchromophyceae exhibit a range of plastid features, including the cryptic non-photosynthetic plastids recently discovered in *Leukarachnion* sp. PRA-24 (Barcytė et al. 2024).

Notably, the branching order within the Synchromophyceae resolved by our phylogenomic analysis, with *Chlamydomyxa* and *Leukarachnion* sp. PRA-24 as a clade excluding *Synchroma*, is consistent with ultrastructural characteristics such as the structure of the cyst wall (Pearlmutter and Timpano 1984; Grant et al. 2009; Barcytė et al. 2024). The depth of divergence between the three major constituent lineages of the Synchromophyceae in both the phylogenomic analysis (fig. 1) and the 18S rRNA gene tree (fig. 6) seems to be best formalized by classifying them as separate orders, two extant (Synchromales and Chlamydomyxales) and one to be established for the clade containing *Leukarachnion* sp. PRA-24 and its uncultured relatives (this is being proposed in a parallel study, in which *Leukarachnion* sp. PRA-24 is redescribed as *Leucomyxa plasmidifera* gen. et sp. nov. and the order Leucomyxales is proposed for it and its uncultured relatives (Barcytė et al. 2024). Interestingly, the existence of a fourth synchromophyte order is indicated by environmental 18S rDNA amplicons from freshwater samples (fig. 6; fig. S1). However, we leave the development of a detailed internal classification of the class for the future, when additional amoeboid algae generally reminiscent of synchromophytes, such as the genera *Myxochloris* or *Rhizochloris* (Pascher 1938), are studied by molecular tools.

#### Perspectives

Here we provide new data that, for the first time, robustly place several previously neglected lineages in the ochrophyte phylogeny. Furthermore, these data help to solidify the position of the root of ochrophytes, dividing the group into two principal clades Chrysista and Diatomista. Our work thus provides a well-resolved framework for interpreting trait evolution in this extremely diverse and ecologically important group. The transcriptome assemblies generated for this study have substantially expanded genome-scale sequence resources for ochrophytes and will be instrumental in future comparative studies aimed at tracing the evolution of specific gene families, as well as the global dynamics of evolution of the gene repertoire in ochrophytes.

However, additional work could still improve the representation of the ochrophyte phylogenetic diversity in phylogenomic datasets. The reconstruction of the phylogenetic backbone of the PX clade will remain incomplete until transcriptome or genome data from *Aurearena cruciata* (a sole representative of the class Aurearenophyceae) and the *incertae sedis* genus *Nematochrysis* are generated and added to the analysis (work in progress). Furthermore, ten additional marine ochrophyte lineages with possible class-level status exist, presently documented only by environmental DNA sequences, including five (MOCH-1 to MOCH-5) recognized before (Massana et al. 2014) and an additional five, here denoted MOCH-6 to MOCH-10, unveiled by our analysis (fig. 6; fig. S1). An additional potentially separate ochrophyte lineage is represented by sequences previously released to databases and assigned to the genus “*Chloromorum*” (fig. 6; fig. S1), which has not been formally described, nor is a culture publicly available. Dedicated culturing efforts, single-cell sequencing, and/or metagenomic approaches will eventually allow for inclusion of the missing lineages into phylogenomic analyses. This will not only illuminate their position and formal taxonomic status among ochrophytes but may also help resolve the remaining uncertainties regarding the deepest ochrophyte relationships, especially the position of eustigmatophytes.

By reevaluating actinophryids as a *bona fide* ochrophyte lineage and showing that *Picophagus flagellatus* lacks any genomic signature for the presence of a plastid, we demonstrate that plastid loss is mechanistically possible in ochrophytes. This suggests that other such cases await discovery. The prime candidate so far, which is lacking genome-scale data, is the dictyochophyte genus *Ciliophrys*, which has no plastid discernible by TEM (Sekiguchi et al. 2002). Furthermore, the *rbcL* sequence previously reported from this organism, which would indicate the presence of a cryptic plastid, was recently revealed to likely result from contamination (Kayama et al. 2020). The existence of aplastidic free-living ochrophytes has implications beyond this group, underscoring the possibility of plastid loss outside the context of reductive evolution associated with parasitism. So far, the only free-living lineage that could be considered a strong candidate for plastid loss is Picozoa, a group of tiny marine flagellates nested among the primary plastid-containing supergroup Archaeplastida (Schön et al. 2021). On the other hand, recurrent plastid loss features prominently in the chromalveolate hypothesis (in any of its variants; Cavalier-Smith 1999; Cavalier-Smith 2018). While the central tenet of the hypothesis is unlikely to be true, our results provide a new impetus to investigate the possibility that some of the aplastidic taxa, specifically related to algal groups with the “chromalveolate plastid”, may represent additional instances of plastid loss.

### Taxonomic Treatments

#### Synchromophyceae S.Horn & C.Wilhelm, emend. D.Barcytė & M.Eliáš

Emended description: Eukaryotic algae with complex plastids, with chlorophylls *a* and *c* or secondarily without photosynthetic pigments. Vegetative stage amoeboid, with the capability to form multinucleate meroplasmodia, without flagella. Cysts or cyst-like wall-covered stage, present, too; flagellated cells may be produced transiently as a putative dispersal stage. Defined as a monophyletic group including *Synchroma grande* R.Schnetter and *Chlamydomyxa labyrinthuloides* W.Archer, but not *Ochromonas triangulata* Vysotskii. Included genera: *Synchroma* R.Schnetter, *Chrysopodocystis* C.Billard, *Guanchochroma* R. Schnetter & M.Schmidt, *Chlamydomyxa* W.Archer, *Leucomyxa* D.Barcytė & M.Eliáš (a new genus being proposed to accommodate the organism so far known as *Leukarachnion* sp. PRA-24; Barcytė et al. 2024).

Note: the class contains the orders Synchromales R.Schnetter & Ehlers (with the sole family Synchomophyceae R.Schnetter & Ehlers, but perhaps to be extended to embrace also the *incertae sedis* genera *Chrysopodocystis* and *Guanchochroma*) and Chlamydomyxales Archer (with the sole family Chlamydomyxaceae Engler). Another order (Leucomyxales) is being proposed elsewhere (Barcytė et al. 2024) to accommodate *Leukarachnion* sp. PRA-24.

#### Picophagea Cavalier-Smith, emend. T.Pánek & M.Eliáš

Emended diagnosis: Biciliate heterokont non-photosynthetic phagotrophic zooflagellates without plastids; lacking stomatocysts. Distinguished from other similar taxa by its unique phylogenetic position defined by molecular phylogenetic analyses, corresponding to a separate phylogenetic lineage closely related to, but distinct from, a clade containing the classes Chrysophyceae and Synchromophyceae).

Note: presently monotypic, the sole species *Picophagus flagellatus* Guillou & Chrétiennot-Dinet in the family Picophagaceae Cavalier-Smith, order Picophagales Cavalier-Smith.

## Methods

### Material sources and culturing

Cultures were purchased from culture collections based on availability, derived from the previous work of the authors involved in this study, or obtained from collaborators (details in table S1). The cultures were grown in 250mL glass flasks, mostly under the culturing conditions recommended by the respective culture collections. Culture growth was monitored by using a fluorometer and once each culture reached mid-exponential phase, cells were harvested over a 2-micron 25-millimeter polycarbonate filter, flash frozen in liquid nitrogen, and stored in a −80° C freezer until RNA was isolated. *Picophagus flagellatus* RCC22 was grown in 3% LB medium (in artificial sea water), *Olisthodiscus luteus* strain K-0444 was cultivated as previously described (Barcytė et al. 2021), and *Chrysoparadoxa australica* EC13 was grown in K-enriched seawater (Wetherbee et al. 2018).

### RNA isolation, sequencing, and assembly

For a majority of strains, RNA was extracted using Machery Nagel plant RNA kit following manufacturers protocol except for cell lysis. This was carried out by bead beating for 5 minutes using a mixture of 0.1 and 0.5mm zirconia/silica beads (BioSpec) in a BioSpec Mini-Beadbeater. Sequencing libraries were constructed using the NEB Next Ultra RNA prep kit and sequenced on the Illumina HiSeq (4000) in paired-end mode (2×150bp). Adapters and low-quality regions were removed using Trimmomatic (Bolger et al. 2014). Trimmed reads were assembled using rnaSPAdes v3.13 (Bushmanova et al. 2018). In the case of *O. luteus* K-0444 and *P. flagellatus* RCC22, total RNA was extracted with TRI Reagent® (TR 118) (Molecular Research Center, Inc., Cincinnati, USA), following standard procedures. Sequencing libraries were prepared by Macrogen Inc. (Seoul, South Korea) using TruSeq Stranded mRNA LT Sample Prep Kit and transcriptome sequencing was performed with the Illumina NovaSeq 6000 platform in pair-end mode (2×151bp and 2×101bp for *O. luteus* and *P. flagellatus*, respectively). *De novo* transcriptome assemblies for *O. luteus* and *P. flagellatus* were obtained using Trinity v2.1.1 (Grabherr et al. 2011). RNA of *C. australica* EC13 was extracted with a Purelink Plant RNA kit (ThermoFisher). Sequencing libraries were prepared and sequenced on Illumina HiSeq 2500 (Novogene, Hong Kong) and assembled with Trinity 2.4.0. Transcripts were translated to protein sequences using TransDecoder (https://github.com/TransDecoder/). WinstonCleaner (https://github.com/kolecko007/WinstonCleaner) was used to identify and remove lowly expressed transcripts and cross-contamination from the assembled transcriptomes. Completeness of the genome or all transcriptomes generated as part of this study were assessed using BUSCO v. 5.5.0 (Manni et al. 2021) with the Stramenopile gene set (table S1).

### Phylogenetic analysis of the 18S rRNA gene

The analysis was carried out with two objectives in mind: (1) to document the position of the ochrophytes (particular strains) included in the phylogenomic analysis in the context of a more densely sampled phylogeny and to confirm their taxonomic identification; (2) to provide a comprehensive picture of the phylogenetic diversity of ochrophytes at the level of major extant lineages, considering also the fact that a vast resource of environmental 18S rRNA gene sequences exists providing a foray into the diversity of uncultured ochrophytes. To the first end, 18S rDNA/rRNA sequences were gathered for the 81 ochrophytes selected for the phylogenomic analysis from various sources. When available and sufficiently complete and accurate, a previously published PCR-derived sequence from the respective strain (or its replica in another culture collection) was used. In the case of the strains where such sequences were unavailable in GenBank or were found two short or chimeric, the respective sequence was extracted from the existing genome or transcriptome assembly and if needed, manually assembled from two or more overlapping contigs. A full 18S rRNA gene sequence assembly was not possible in some cases, so only a partial sequence (rather than a longer, potentially inaccurate one) was accepted for the analysis. In a few cases the reference 18S rRNA gene sequence was obtained for the respective organisms from the metadata accompanying the protein sequence database EukProt3 (Richter et al. 2022). No 18S rRNA gene sequence could be found for one of the targeted ochrophytes (Chrysophyceae sp. TOSAG23-5, represented by a single-cell amplified genome). Details on the reference sequences with additional comments are provided in table S1.

These sequences were combined with 180 additional 18S rRNA sequences that were selected based on previously published phylogenies of ochrophytes or stramenopiles and our own exploration of sequences in GenBank. Briefly, blastn searches of the NCBI nr nucleotide sequence database were carried out with different regions of selected ochrophyte 18S rRNA gene sequences as queries (to maximize the chance of finding even partial sequences representing phylogenetically unique lineages). The best hits were combined and exploratory trees were inferred with the FastTree algorithm available at the ETE3 server (https://www.genome.jp/tools-bin/ete). Sequences representing different major clade were evaluated for their length and possible chimeric origin, and a representative subset of them was retained, prioritizing longer sequences and aiming to sample more or less evenly across the ochrophyte radiation and comprehensively in terms of the established major taxa (classes, orders), lineages *incertae sedis*, and clades comprised solely of environmental DNA sequences. Given the special interest in Synchromophyceae we also included two sequences derived from metagenome assemblies and previously shown to be part of the “*Leucomyxa* clade” (i.e. related to the organism currently referred to as *Leukarachnion* sp. PRA-24; Barcytė et al. 2024). To properly represent the phylogenetic span of Bolidophyceae, we also included the 18S rRNA gene sequences provided in a supplement to a report on the genome sequencing of several Bolidophyceae members (Ban et al. 2023). Non-ochrophyte stramenopiles were sampled less comprehensively, primarily including representative sequences from taxa and eDNA clades from previous analyses known to be phylogenetically closest to ochrophytes. The names of the taxa (genera, species) represented by the sequences were updated based on the recent taxonomic revisions and thus not always correspond to the names used in the respective GenBank records.

A multiple alignment of the final set of 260 sequences was constructed with MAFFT version 7 (Katoh and Standley 2013) using the L-INS-i method, inspected by eye and trimmed manually to remove poorly conserved positions (with 1,820 positions kept in the final alignment). A ML tree was inferred from the alignment with IQ-TREE multicore version 2.2.5 (Minh et al. 2020), the substitution model TN+F+I+R7 automatically selected by the program as best fitting the data, and ultrafast bootstrap approximation with 1,000 bootstrap replicates. For presentation purposes the tree was processed with the aid of iTOL (Letunic and Bork 2021).

### Phylogenomic dataset construction

The phylogenomic datasets were built with the aid scripts that are part of the PhyloFisher package (Tice et al. 2021; Jones et al. 2024). Putative orthologs were collected from the predicted protein sets by employing the fisher.py script using the default settings and utilizing the diatom *Phaeodactylum tricornutum* as a blast seed. The retrieved sequences were added to the alignments in PhyloFisher’s starting database using working_dataset_constructor.py with default settings. Single gene datasets were aligned using MAFFT (--globalpair --maxiterate 1000), alignment uncertainty and errors were filtered using DIVVIER (--partial -mincol 4 -divvygap) (Ali et al. 2019) and single gene trees were inspected for paralogy and contamination using Parasorter and forest.py. Once orthologs for all taxa were identified, final taxon selection was carried out resulting in a final dataset of 240 orthologous amino acid sequences and 112 taxa (Stram dataset) or 94 taxa (Ochro dataset) was created using select_taxa.py, select_orthologs.py, and prep_final_dataset.py. The final concatenated dataset was generated using matrix_constructor.py. Our final datasets were once again filtered and aligned as described above. Single gene alignments were concatenated into a super matrix, and a maximum likelihood (ML) tree was inferred using IQ-TREE v 1.6.7 (Nguyen et al. 2015) under the LG+C60+G4 mixture model. The resulting ML tree was used as a guide tree under the same model to estimate the PMSF profiles (Wang et al. 2018). Support values were then estimated using 500 nonparametric bootstrap replicates under the LG+C60+G4+PMSF model. Bayesian analyses were inferred using PhyloBayes-MPI v. 1.5 (Lartillot et al. 2013) with the CAT+GTR+G4 model. Four independent Markov Chain Monte Carlo (MCMC) chains were run for >10,000 generations (all sampled, burnin=20%).

For the fast site removal (FSR) analysis, per site evolutionary rates were estimated and sites in the alignment were sorted from fastest to slowest evolving utilizing fast_site_remover.py. This was followed by sequential removal of 3,000 fastest sites at a time generating increasingly smaller datasets at each step. Similarly, for the heterotacheous sites removal (HSR), the most heterotacheous sites were removed in a stepwise fashion (3,000 sites at a time) using heterotachy.py, producing iteratively smaller datasets until no further sites could be removed. Random subsampling (RGS) of 20%, 40%, 60%, or 80% of genes that comprise the complete phylogenomic dataset was carried out using random_resampler.py. For each of the FSR, HSR, and RGS analyses, ML trees were reconstructed under the LG+G+C20-PMSF model with 100 nonparametric bootstraps for each step or subsample. Additionally, the AU test (implemented in IQ-TREE) was conducted on the ML trees, constrained topologies of interest (inferred using IQ-TREE under the LG+C60+G4 mixture model model), and 100 distinct local topologies saved during the initial ML analysis (-wt option in IQ-TREE).

### Sequencing of the *Picophagus flagellatus* RCC22 genome

DNA extraction was performed from pelleted cells (together with the co-cultured prokaryotes) from a modified protocol originally used for plants (Dellaporta et al. 1983). Modifications included additional steps of RNAse (RNAse H; the final concentration 0.1 mg/ml) and proteinase treatments (proteinase K; the final concentration 0.2 mg/ml) followed by phenol-chloroform extraction before the final DNA precipitation. The sequencing library was prepared by Macrogen Inc. (Seoul, South Korea) using TruSeq DNA PCR-Free Protocol (Illumina, San Diego, CA) and genome sequencing was performed with the Illumina HiSeq 2000 platform in pair-end mode (2×151bp). Raw reads were trimmed by Trimmomatic v. 0.32 and the genome was *de novo* assembled by SPAdes 3.13.1. (Bankevich et al. 2012) using default parameters under the “careful” mode to minimize number of mismatches in the final contig. The completeness of the genome assembly was evaluated using BUSCO v5.5.0 with the Stramenopile gene set. In addition to scaffolds derived from the nuclear and mitochondrial genome of *P. flagellatus* itself, the final assembly also contains genomes of prey bacteria, so caution needs to be exercised when exploring these data.

### Exploration of the *P. flagellatus* RCC22 transcriptome and genome assemblies

Protein sequences encoded by the plastid genome of *Olisthodiscus luteus* K-0444 (NC_057170.1) and the mitochondrial genome of *Vischeria* sp. CAUP Q 202 (KU501221.1) were used as queries for a tblastn search to identify putative organellar genome sequences in the *P. flagellatus* RCC22 genome assembly. Significant hits were evaluated by blastx searches against the NCBI non-redundant protein sequence database, resulting in the identification of a single scaffold corresponding to the mitochondrial genome, whereas all candidates for possible plastid genome-derived sequences were identified as bacterial contaminants. Homologs of proteins of special interest (table S3 and table S4) were searched primarily in the *P. flagellatus* transcriptome assembly (and proteins inferred from it) and if not identified, in the genome assembly as well. In the case of components of plastid protein import complexes and proteins involved in plastid division, two other non-photosynthetic ochrophytes, *Leukarachnion* sp. PRA-24 (transcriptome assembly; Barcytė et al. 2024) and *Paraphysomonas imperforata* CCMP1604 (annotated genome assembly; https://phycocosm.jgi.doe.gov/Parimp1_4/Parimp1_4.home.html) were used as a reference. The searches were performed using blast with reference sequences selected based on literature data as queries, or (for Omp85, Tic20, Tic22, Tic110, Tom40, Sam50, Tim17/22/23) using HMMER3.1 searches (Eddy 2011) with profile HMMs created from seed alignments corresponding to relevant protein families as defined in the Pfam database (Mistry et al. 2021). Hits were evaluated by blastp searches against the NCBI nr database, and when necessary by HHpred (https://toolkit.tuebingen.mpg.de/tools/hhpred; Zimmermann et al. 2018) or by simple phylogenetic analyses using FastTree implemented in the ETE3 server (https://www.genome.jp/tools-bin/ete) and a set of best blastp hits obtained from the EukProt3 database. Proteins inferred from the *P. flagellatus* transcriptome assembly were mapped onto the metabolic pathways in the KEGG database using BlastKOALA (Kanehisa et al. 2016) and sequences assigned to enzymes of pathways of specific interest were evaluated to ascertain whether they belong to *P. flagellatus* or bacteria contaminating the culture. To evaluate the subcellular localization of the two enzyme homologs of the lysine biosynthesis pathway, TargetP 2.0 (“non-plant” setting; https://services.healthtech.dtu.dk/service.php?TargetP-2.0; (Almagro Armenteros et al. 2019), Predotar (https://urgi.versailles.inra.fr/predotar/; Small et al. 2004), PredSL (set to “non-plant sequences”; http://aias.biol.uoa.gr/PredSL/input.html; Petsalaki et al. 2006), PrediSi (http://www.predisi.de/; Hiller et al. 2004), and DeepLoc 2.0 (Thumuluri et al. 2022) were used. The former three tools provided also prediction of possible mitochondrial targeting of the proteins, which was additionally tested using MitoFates (arbitrarily with the default setting to “fungi”; http://mitf.cbrc.jp/MitoFates/cgi-bin/top.cgi; Fukasawa et al. 2015). Detailed phylogenetic analyses were carried out for selected *P. flagellatus* proteins, using sets of homologous sequences collected with blast from the NCBI nr database, EukProt3, and taxon specific resources (transcriptome assemblies from *A. sol*, *Leukarachnion* sp. PRA-24, and ochrophytes sequenced in this study). To obtain representative prokaryotic outgroups, strategically selected sequences were used as a query for PSI-BLAST search in clustered NCBI nr database (nr_pro70) available at toolkit.tuebingen.mpg.de/tools/psiblast (Zimmermann et al. 2018). Protein sequences were aligned with MAFFT, using L-INS-I method with BLOSUM 30. Alignments were trimmed manually. Maximum likelihood phylogenetic analyses were performed with RAxML version 8.2.12 under the LG4X substitution model (Stamatakis 2014). Branch supports were estimated by rapid bootstrapping (N = 100).

## Acknowledgements

We thank Karin Jaške, Tatiana Yurchenko, and Martin Sokol for their assistance with obtaining sequence data from *O. luteus* and *P. flagellatus*. This work was supported by the National Science Foundation DEB Grant #1541510 to CL, the Czech Science Foundation (project 23-06203S to T.P. and M.E) and the European Union under the LERCO project number CZ.10.03.01/00/22_003/0000003 via the Operational Programme Just Transition. Ma.K. by the ERD fund “Centre for Research of Pathogenicity and Virulence of Parasites” (no. CZ.02.1.01/0.0/0.0/16_019/0000759) and by Czech Science Foundation (project #: 22-22538S). E.D.S. was supported in part by the MSCA-IF SMART (CZ.02.2.69/0.0/0.0/20_079/0017809). This research was conducted in part using computational resources and services at the Center for Computation and Visualization, Brown University. Computational resources were provided in part by the Rhode Island Consortium for Coastal Ecology Assessment, Innovation and Modeling which is funded in part by the National Science Foundation under EPSCoR Research Infrastructure Improvement (Award #OIA-1655221) and by IT4Innovations National Super Computer Center, TUO, Ostrava, Czech Republic (projects #OPEN-23-29). A Grant in Aid of Research from the Phycological Society of America was also awarded to KT in support of this work.

## Data availability

Transcriptome and genome assemblies, alignments, and trees reported in the paper are available from Figshare######.

---

**Figure S1:** Ochrophyte phylogeny inferred from sequences of the 18S rRNA gene. The tree was inferred from an alignment of 260 sequences (1,820 positions after trimming) with IQ-TREE and ultrafast bootstrap approximation with 1000 bootstrap replicates (bootstrap values are shown at the corresponding branches when ≥50%). The root of the tree was placed between the three representatives of the stramenopile clade Bigyra and the remaining ochrophytes (all belonging to the clade known as Gyrista; see also fig. S2). Major clades, corresponding to formally established taxa (mostly classes) and informally denoted eDNA groups (MAST, MOCH, and the “Abyssal” clade), were annotated based on the literature, with five newly recognized ochrophyte clades (highlighted in red) solely consisting of sequences from environmental surveys of marine habitats designed as MOCH-6 to MOCH-10 (“MOCH” = “marine ochrophytes”) as an increment to the MOCH-1 to MOCH-5 clades defined previously (Massana et al. 2014). Sequences corresponding to strains (or vouchers) included in the phylogenomic analysis are highlighted with a turquoise background.

**Figure S2:** Maximum likelihood phylogeny of stramenopiles recovered from 112 taxa, 240 genes and 103,167 sites (the Stram dataset). The tree root is placed arbitrarily between the two principal stramenopile assemblages (Bigyra and Gyrista) suggested by previous analyses including a non-stramenopile outgroup (Derelle et al. 2016). Non-parametric PMSF bootstrap support (BS) values (n=500) and PhyloBayes posterior probabilities (PP) are shown on the branches as follows: BS/PP. Branches with black circles received maximum support in both analyses. Ochrophytes newly sequenced in this study are bolded and colored blue.

**Figure S3:** Effects of random gene subsampling on both our Stram and Ochro datasets. Box-and-whisker plots showing support for randomly sampled subsets of genes from the Ochro and Stram datasets (Ochro shown on top and Stram on the bottom). Each subsampled percentage is its own plot with non-parametric bootstrap support (n=100) for on the x-axis and relationships being tested on the y-axis. The number of individual datasets required to sample every gene at 95% probability is shown above each plot.

**Figure S4:** Maximum likelihood phylogeny of stramenopiles in the broader context of SAR (Act dataset) recovered from 75 taxa, 240 genes and 95,605 sites, with Rhizaria and Alveolates as outgroup taxa. Non-parametric PMSF bootstrap support (BS) values (n=500) and Phylobayes posterior probabilities (PP) are shown on the branches as follows: BS/PP. Branches with black circles received maximum support in both analyses.

**Figure S5.** Assessment of the completeness of the genome assembly for *Picophagus flagellatus* RCC22 using BUSCO v5.5.0 (using the Stramenopile gene set). For comparison, genome assemblies of three additional ochrophytes, considered to be essentially complete, were included. The sources of these assemblies were as follows: *Ectocarpus* sp. Ec32 – https://phycocosm.jgi.doe.gov/Ectsil1/Ectsil1.info.html; *Phaeodactylum tricornutum* – https://www.ncbi.nlm.nih.gov/datasets/genome/GCF_000150955.2/; *Vischeria* sp. C74 – (https://figshare.com/s/c4bf156c2764ba410c30.

**Figure S6**. Phylogenetic analysis of organellar and cytoplasmic aspartyl-tRNA synthetases showing the position of proteins from *Picophagus flagellatus* and *Actinophrys sol* within a clade of dually-targeted enzymes of plastid origin and lack of Ochrophyta homologs with mitochondrial origin. A selection of asparaginyl-tRNA synthetases was used as an outgroup. Only bootstrap values ≥50 are shown. Clades not of interest here were collapsed as closed triangles and the tree was rooted arbitrarily. The clade of mitochondrial homologs that branched within the collapsed clade is shown in detail separately. Multiple sequence alignment contained 308 protein sequences and 431 positions after trimming, it is available in Figshare link ####.

**Figure S7.** Broad scale phylogenetic analysis of diaminopimelate decarboxylase (DAPDC). Only bootstrap values ≥50 are shown. The tree is arbitrarily rooted. Multiple sequence alignment contained 182 protein sequences and 416 positions after trimming, it is available in Figshare link ####.

**Figure S8.** Phylogenetic analysis of well-supported clade of diaminopimelate decarboxylase (DAPDC) that includes ochrophyte homologs (for context see Figure S7). The trees were inferred with the ML method in RAxML v8.2.12. (using the LG4X substitution model). Branch support was assessed by rapid bootstrapping (N=100). Only bootstrap values ≥50 are shown. The tree is arbitrarily rooted. Multiple sequence alignment contained 133 protein sequences and 412 positions after trimming, it is available in Figshare link ####.

**Table S1:** Organisms used in study

Source, data type, and updated nomenclature and taxonomy for data used in this study. all transcriptomes or genomes included as part of this study were assessed using BUSCO v. 5.5.0 data were analyzed using either the stramenopile, alveolata_odb10 database, or the Eukaryota_odb10 database depending on lineage.

**Table S2:** Trees in Newick format from phylogenomic analyses.

**Table S3**: Proteins implicated in plastid biogenesis and reproduction identified in selected non-photosynthetic ochrophytes but not *Picophagus flagellatus* RCC22.

**Table S4:** Nucleus-encoded proteins of special interest from *Picophagus flagellatus* RCC022.

