## Supplementary figures and images for "Multiple plastid losses within photosynthetic stramenopiles revealed by comprehensive phylogenomics"

### Supplemental Figures

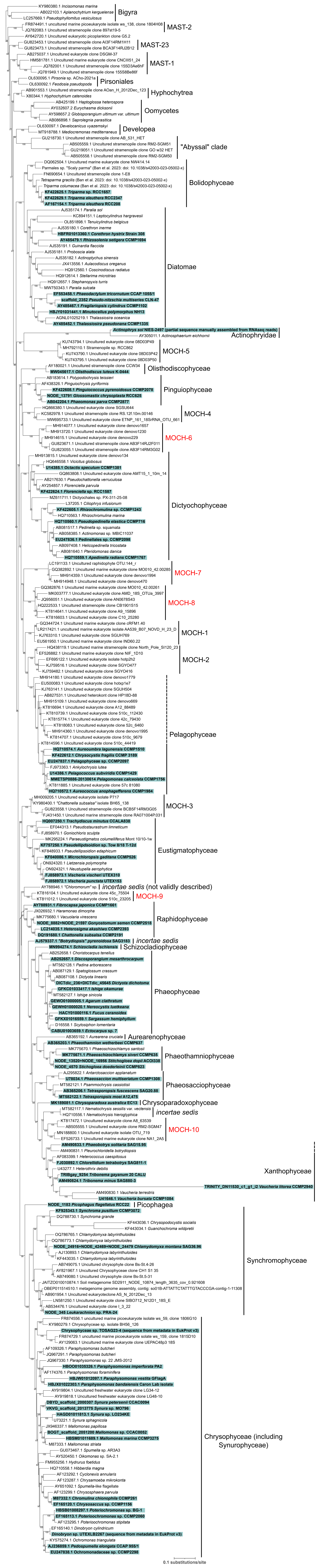

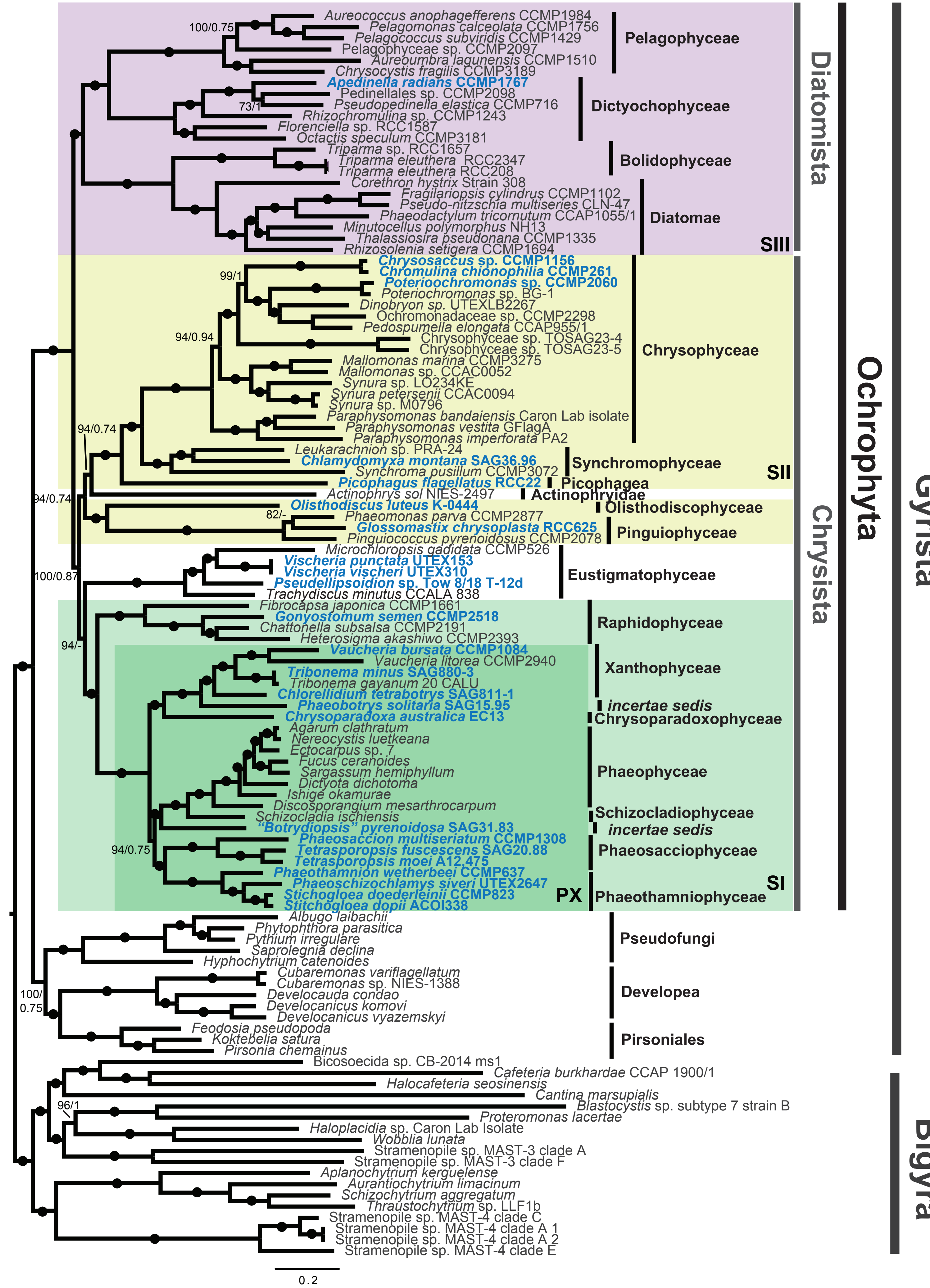

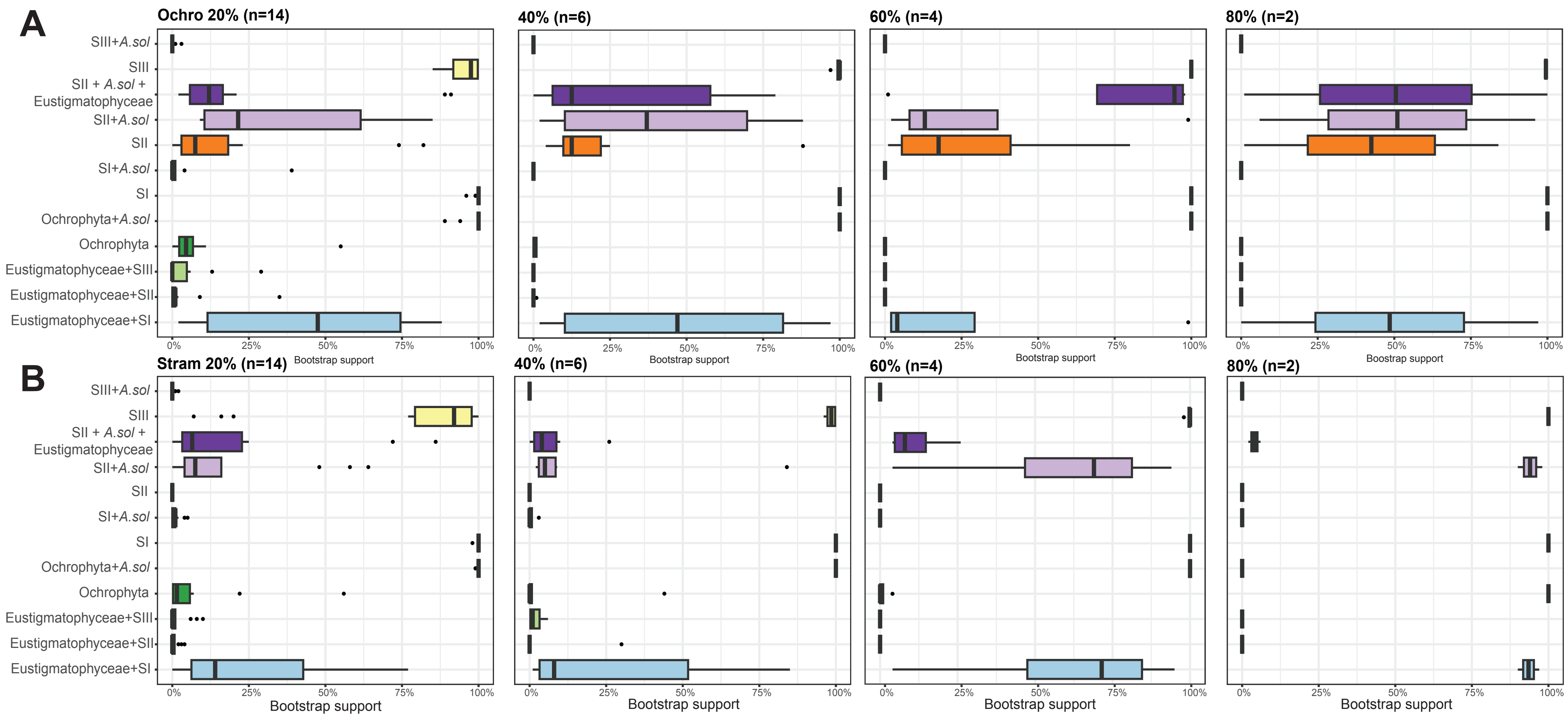

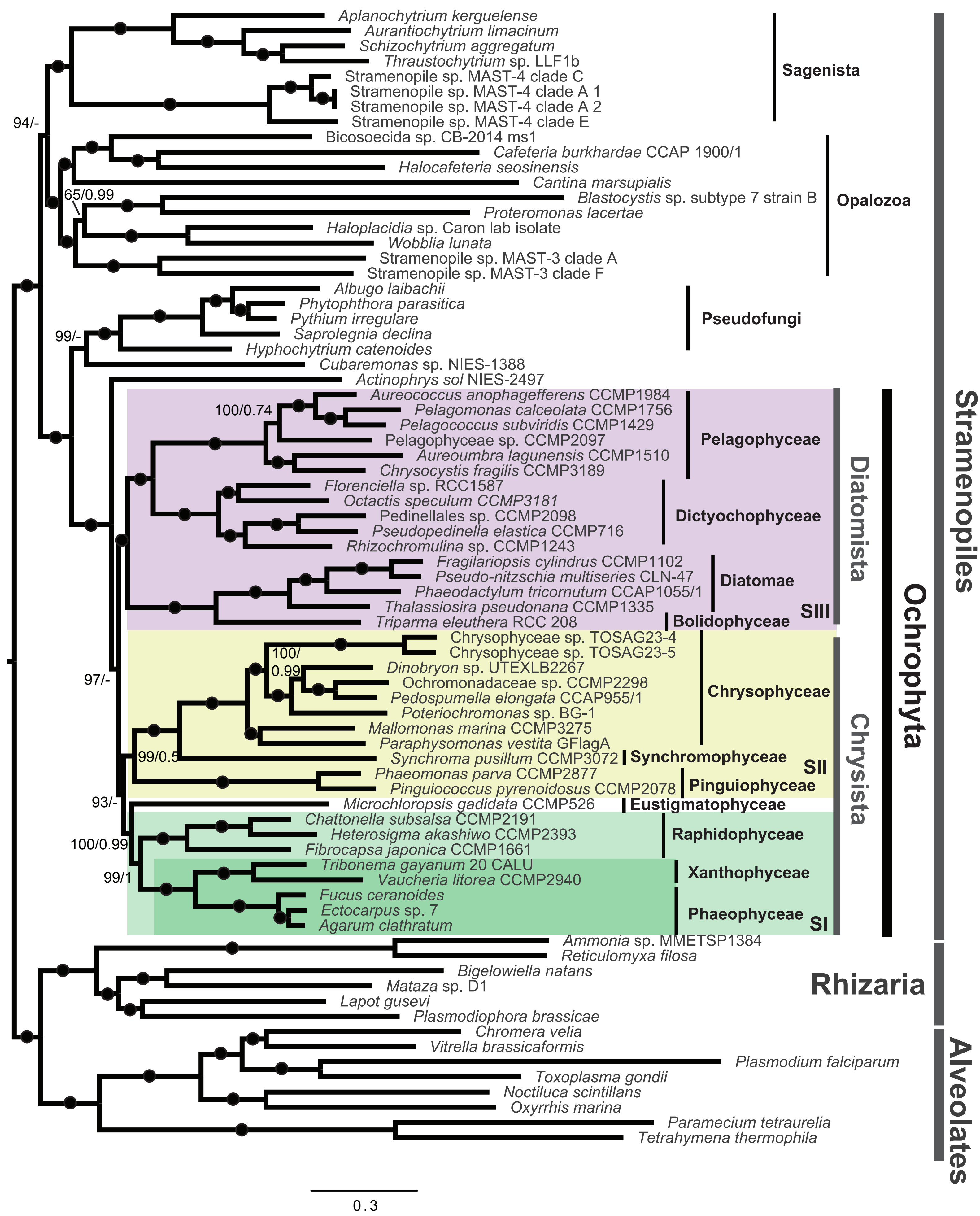

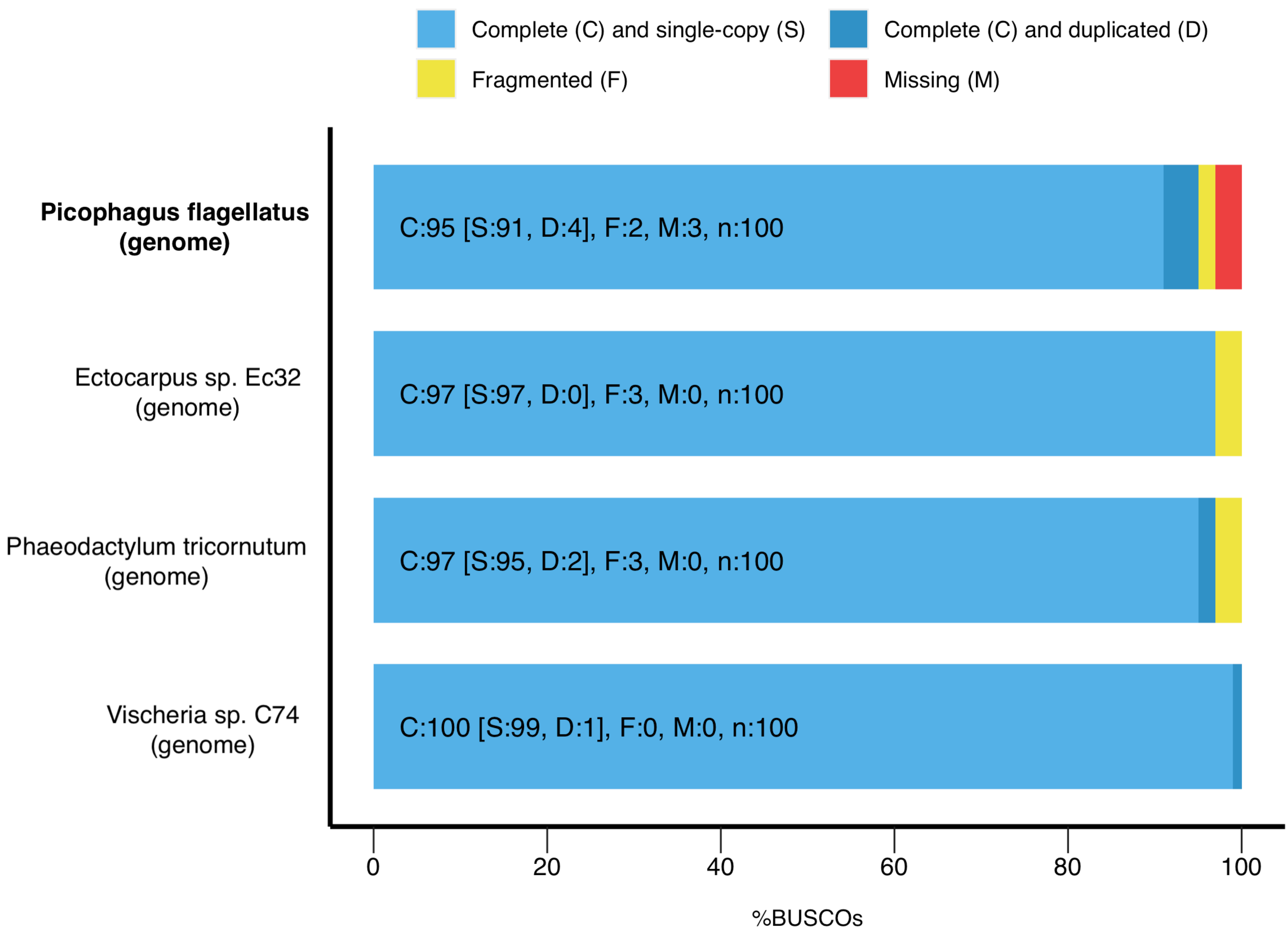

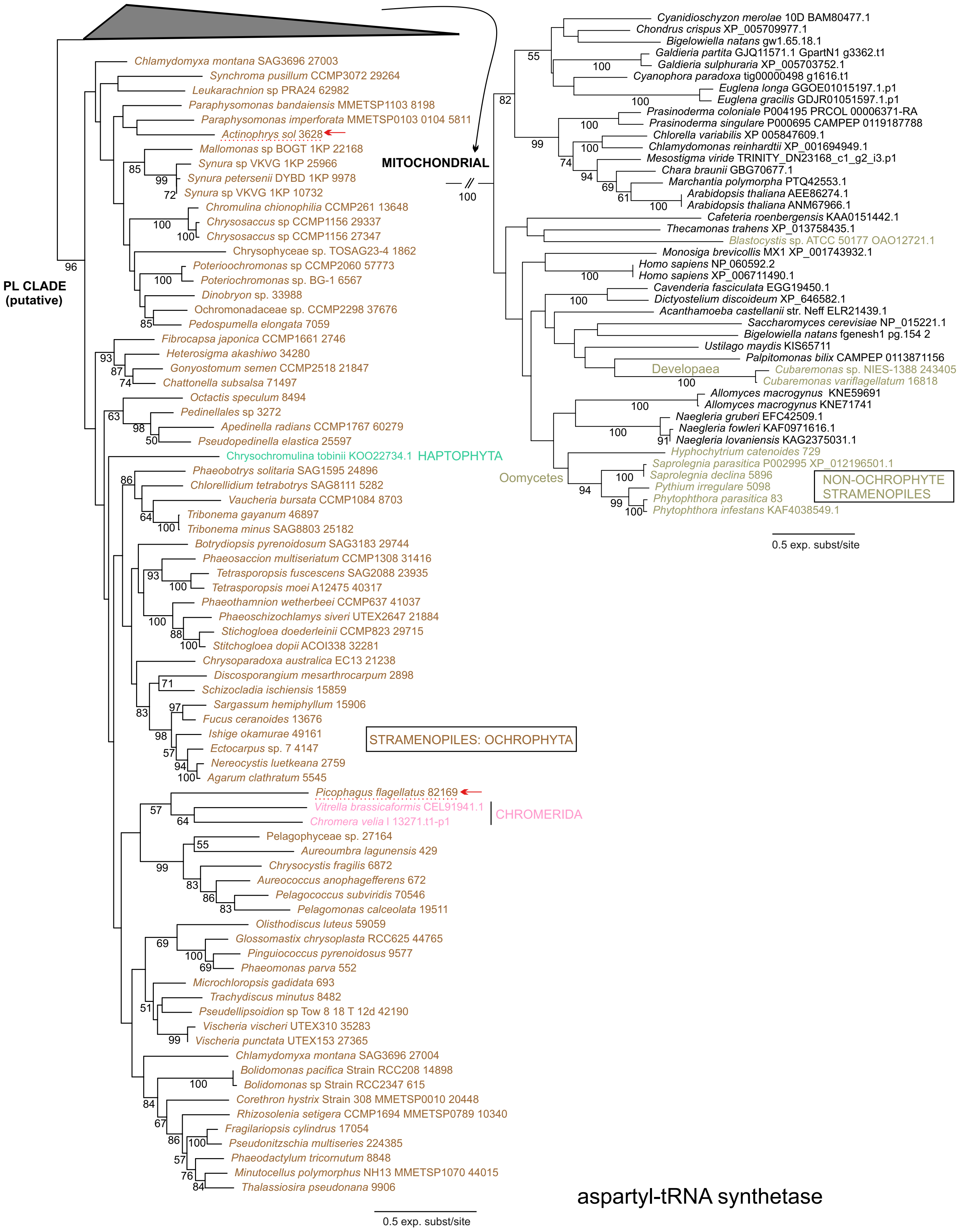

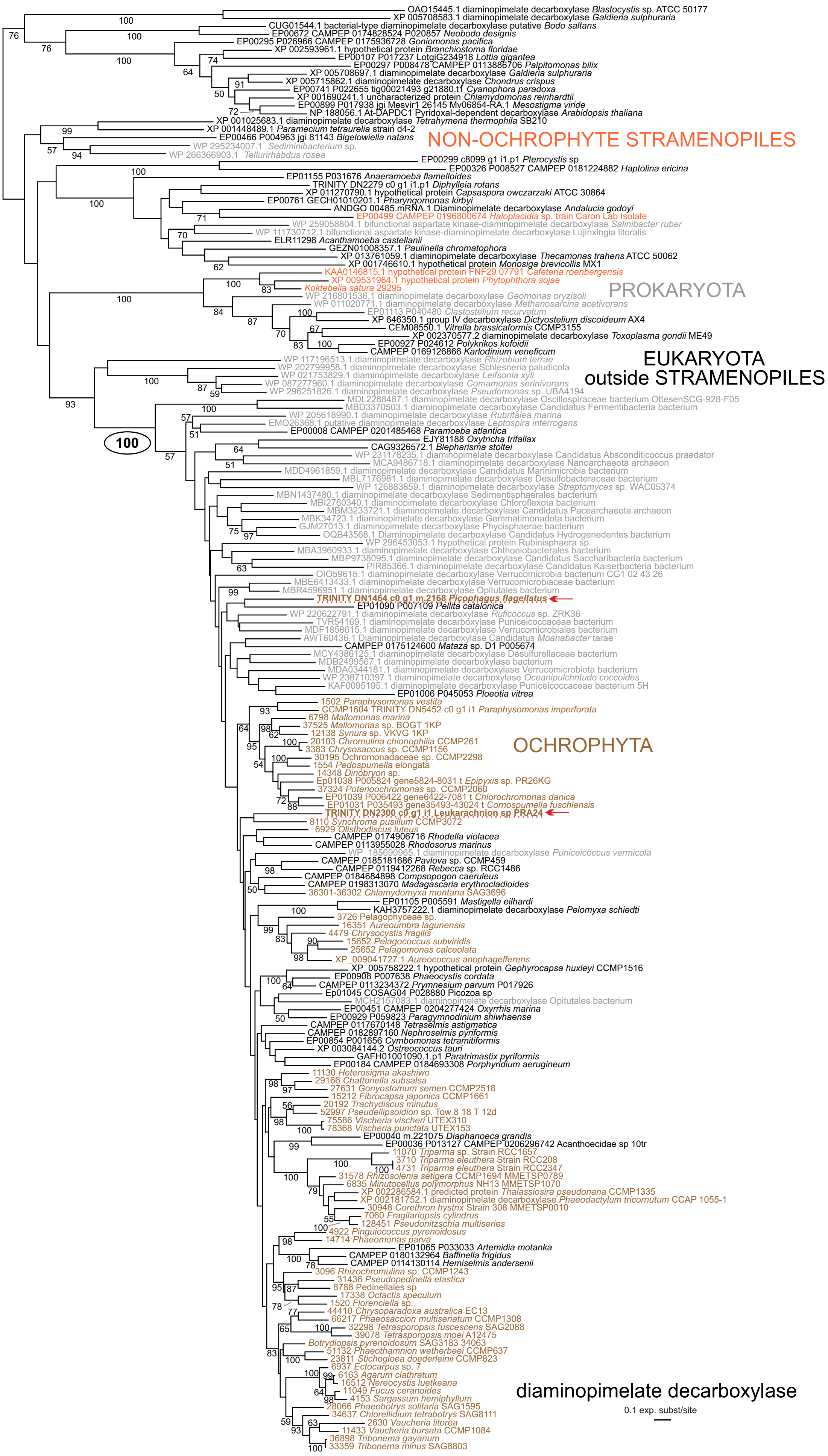

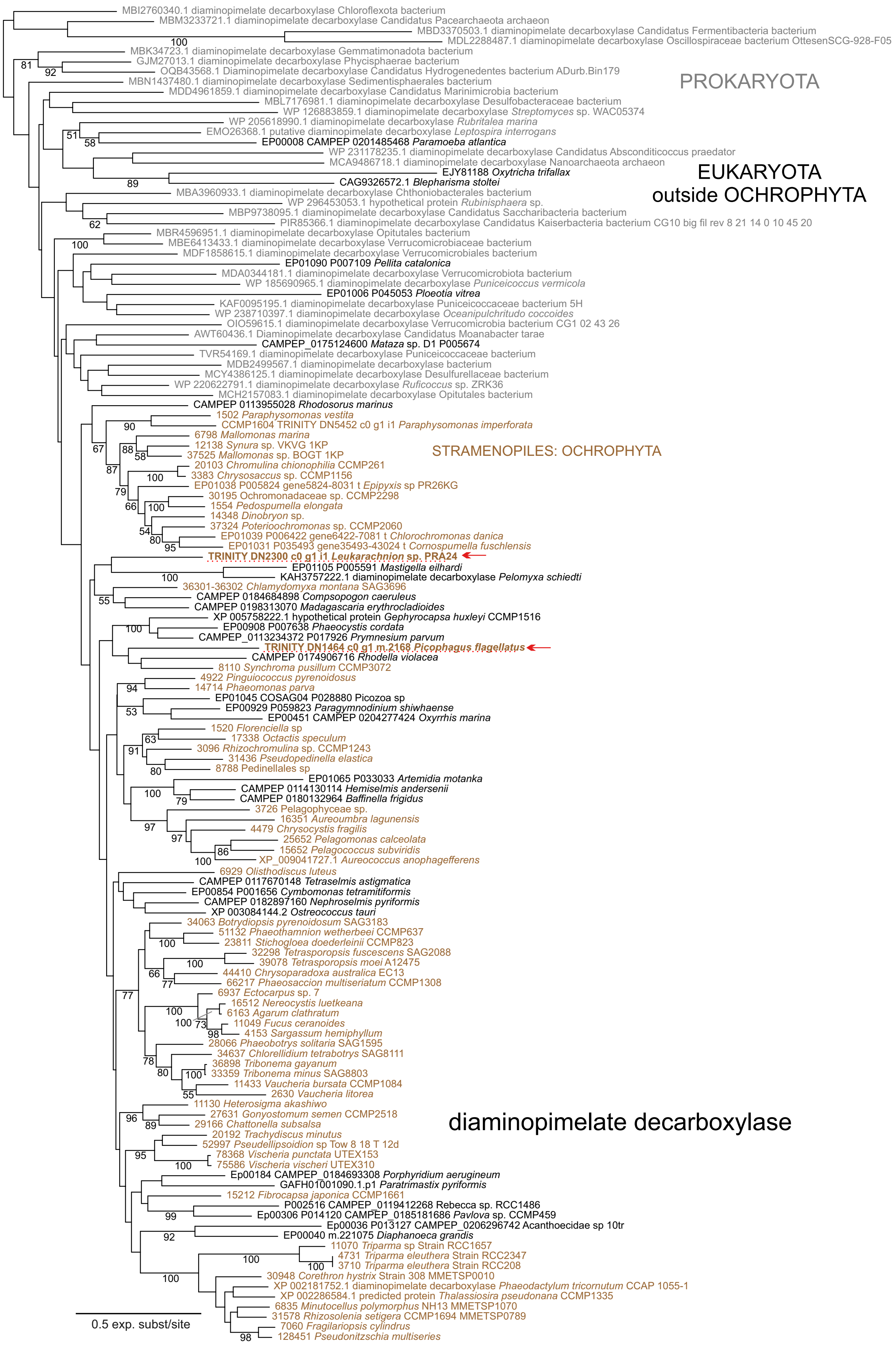
